# Predicting lapses of attention with sleep-like slow waves

**DOI:** 10.1101/2020.06.23.166991

**Authors:** Thomas Andrillon, Angus Burns, Teigane MacKay, Jennifer Windt, Naotsugu Tsuchiya

**Author notes:** Corresponding author: Thomas Andrillon. Postal Address: Monash Biomedical Imaging, 770 Blackburn Road, Clayton 3168, Victoria, Australia. Data and Code Availability: Raw data: https://osf.io/ey3ca/?view_only=680c39e7065649c3b783a4efec0a1a94 Code used for analyses: https://github.com/andrillon/wanderIM.

## Abstract

Attentional lapses are ubiquitous and can negatively impact performance. They correlate with mind wandering, or thoughts that are unrelated to ongoing tasks and environmental demands. In other cases, the stream of consciousness itself comes to a halt and the mind goes blank. What is happening in the brain that leads to these mental states? To understand the neural mechanisms underlying attentional lapses, we cross-analysed the behaviour, subjective experience and neural activity of healthy participants performing a task. Random interruptions prompted participants to indicate their mental states as task-focused, mind-wandering or mind-blanking. High-density electroencephalography revealed the occurrence of spatially and temporally localized slow waves, a pattern of neural activity characteristic of the transition toward sleep. These slow waves accompanied behavioural markers of lapses and preceded reports of mind wandering and mind blanking. Furthermore, the location of slow waves distinguished sluggish versus impulsive behaviours, mind wandering versus mind blanking. Our results suggest attentional lapses share a common physiological origin: the emergence of local sleep-like activity within the awake brain.

## Introduction

The human brain sustains the stream of our conscious experiences. Attention can direct cognitive resources toward the external world and enable the selection and amplification of information relevant to an individual’s current behavioural goals^1^. But attention can also turn inward, as is the case when we focus on internally generated task-unrelated thoughts, a phenomenon usually referred to as mind wandering^2^. Recent investigations have also shown that the stream of thoughts can come to a pause, as when individuals who are awake are left with the feeling of an empty mind (mind blanking)^3^.

Shifts of attention towards the internal world, invoking mind blanking and mind wandering, can occur spontaneously without our knowledge or will^4^. In fact, a characteristic feature of attention is its fleeting nature and the difficulty to maintain it on a task for long periods of time^1, 5^. In this paper, we define lapses of attention as the shift of the focus of thoughts away from the task at hand or environmental demands. The consequences of attentional lapses are very diverse. At the behavioural level, they can result in a lack of responsiveness or sluggish reactions, but they can also result in impulsive responses^6^. Curiously, these behavioural failures can be accompanied by a lack of conscious awareness and the absence of mental activity (mind blanking^3^), or rich, spontaneous mental activity (mind wandering^2^).

It is yet unclear whether these different types of attentional lapses (sluggish vs. impulsive behaviours; mind-blanking vs. mind-wandering) belong to a disparate family of behavioural and phenomenological events^7, 8^, each of them associated with different physiological causes,^9, 10^ or whether they can be traced back to common underlying physiological causes^11^. Previous models of mind wandering have proposed that mind wandering and mind blanking might arise in distinct neurophysiological states^3, 9, 10^. However, the fact that both sluggish and impulsive responses increase following sleep deprivation^12, 13^ and in individuals with attentional deficits^6, 14^ implies a common mechanism. Likewise, sleepiness has been associated with both mind wandering and mind blanking^15, 16^ despite these two mental states being phenomenologically distinct^3^. Furthermore, investigations of the sleep onset period (hypnagogia) indicate that subjective experiences resembling mind wandering (focus on internally generated contents) and mind blanking (loss of awareness) can both occur at the border between wakefulness and sleep^17, 18^. Interestingly, these studies seem to associate lapses with pressure for sleep, suggesting an involvement of fatigue.

Indeed, each hour spent awake comes at the cost of mounting sleep pressure. Past research suggests that the need for sleep might only be dissipated by sleep itself,^19^ as sleep plays a vital role in neural homeostasis^20^. When individuals are prevented from sleeping for extended periods of time (as in sleep deprivation studies), a subset of brain regions can start displaying electroencephalographic (EEG) signatures of non-rapid eye-movement (NREM) sleep in the form of sleep-like slow waves (within the delta (1-4 Hz) or theta (4-7 Hz) range), despite individuals being behaviourally and physiologically awake^21, 22^. These sleep-like slow waves within wakefulness are referred to as “local sleep” in contrast with the global whole-brain transition commonly observed at sleep onset^22–25^. Importantly, both local and global transitions toward sleep are characterised by the same neural signature: the occurrence of high-amplitude slow waves.

It has been proposed that the multiplication of sleep-like slow waves could perturb brain functions and cause behavioural lapses^22^. In fact, during sleep, slow waves are associated with episodes of widespread neural silencing^26^, which have been connected to sensory disconnection, behavioural unresponsiveness and the loss of consciousness^27, 28^. Intracranial studies in humans and rodents showed that slow waves during waking are associated with reduced neuronal firing, resulting in behavioural errors^21, 22^. Wake slow waves can also be detected in human non-invasive recordings^29–31^ and here again the amount of slow waves recorded in a given brain region correlates with the number of errors performed in a task recruiting this specific brain region^29, 30^. These results strongly suggest that slow waves could explain the behavioural component of attentional lapses^22^. However, the impact of slow waves on conscious experience, including mind wandering and blanking, is still unclear.

In a recent review, we proposed that local sleep, manifested as the occurrence of slow waves in waking, could not only explain the behavioural consequences of attentional lapses, both regarding sluggish and impulsive responses, but also their phenomenology, like mind wandering or blanking^11^. We also argued that local sleep is not an extreme phenomenon, occurring only when individuals are pushed to their limit, but could occur in well-rested individuals^31^ and explain the occurrence of lapses in our everyday lives. To test this framework, we rely on the detection of slow waves and formulate three different hypotheses as follows: (i) Can slow waves predict, at the single trial level, both sluggish and impulsive behaviours in well-rested individuals? (ii) Are slow waves associated with both mind wandering and blanking? (iii) Does the location of slow waves (i.e. which electrodes show slow waves) differentiate between sluggishness and impulsivity, mind-blanking and mind-wandering? Through these hypotheses we will test the idea that local slow waves could act as a functional switch, transiently perturbing the functioning of a given cortical network. Accordingly, a common physiological event (local slow waves) could lead to drastically different outcomes depending on its location within the brain.

To test these hypotheses, we cross-examined the behavioural performance, subjective reports and physiological data from healthy individuals (N=26) performing an undemanding Go/NoGo tasks^32^ (Figure 1a). We sampled participants’ subjective experience by interrupting them during the task and asking them a series of questions about their mental states prior to the interruption, including whether they were focusing on the task, mind-wandering or mind-blanking (Figure 1b). Finally, we recorded their brain activity using high-density scalp EEG and pupil size as objective proxies for participants’ level of vigilance^33^. Importantly, here participants were neither sleep deprived nor placed in conditions favouring dozing off (i.e. not in bed or reclined, task requiring constant attention and responses).

**Figure 1.**
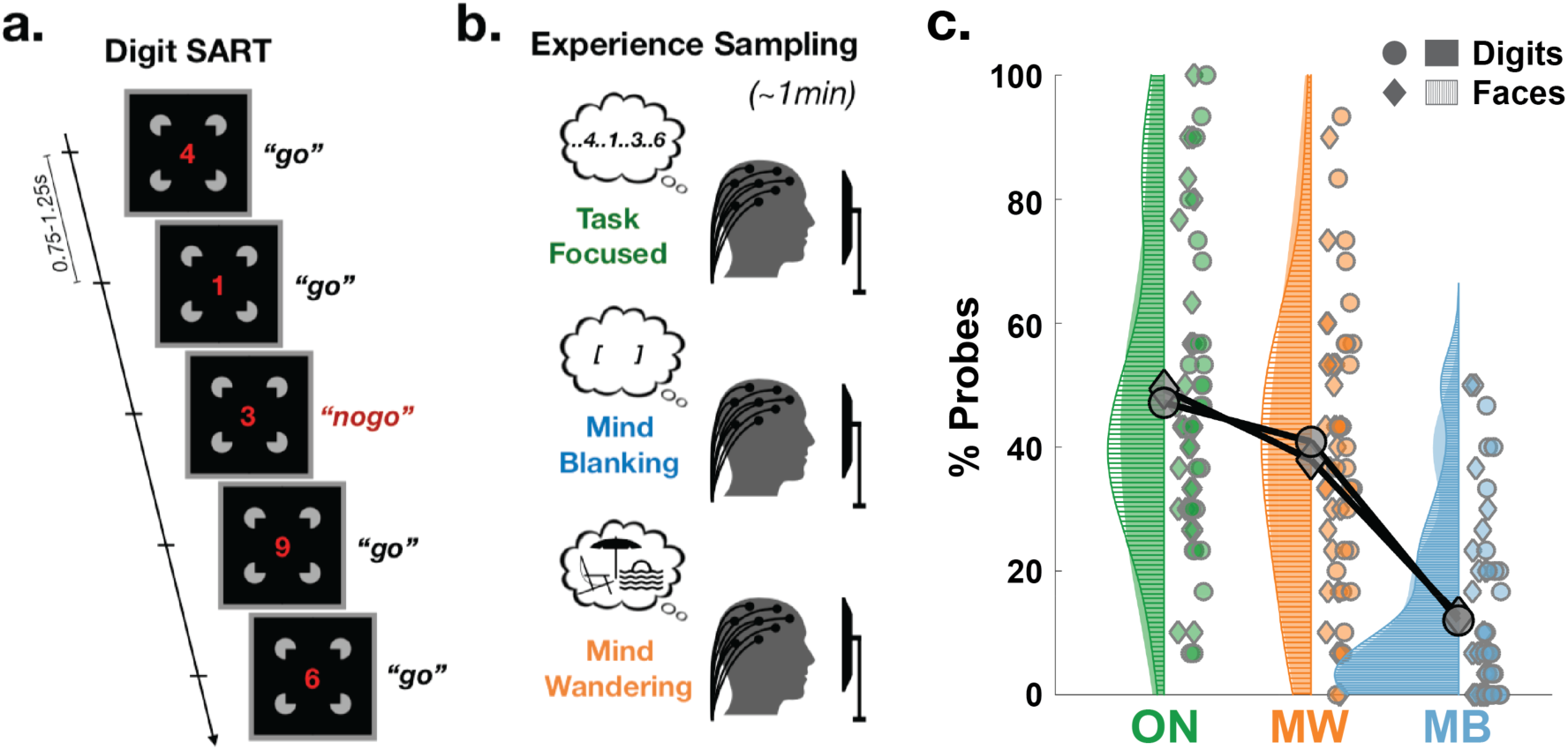
Experimental design and hypotheses. **a:** Participants performed both a SART on faces stimuli (NoGo trials: smiling faces, stimuli shown because of BioRxiv’s guidelines) and a SART on digits (NoGo trials: “3”). Face/Digit presentation was continuous (new face/digit every 0.75-1.25s). **b:** Every 30 to 70s, participants were interrupted and instructed to report their mental state (see Online and Supplementary Methods). Most importantly, they were asked to indicate whether they were focusing on the task (task-focused: ON), thinking about nothing (mind-blanking: MB) or thinking about something other than the task (mind-wandering: MW). High-density EEG and pupil size were continuously recorded throughout the task. **c:** Proportion of mental states reported during probes categorized as task-focused (ON, green), mind-wandering (MW, orange) and mind-blanking (MB, blue) during the tasks with Digits (circles for each individual participant; filled surfaces for smoothed density plot) and Faces (diamonds and surfaces with horizontal stripes). Grey diamonds and circles show the average across participants.

## Results

### Task performance and subjective experience

The Go/NoGo tests (see Online Methods) require participants’ sustained attention, but our participants declared focusing on the task only in ∼48% of the probes (Face Task: 49.4 ± 4.9%; Digit Task: 47.2 ± 5.1%; mean ± Standard Error of the Mean (SEM) across N=26 participants; Figure 1c). The rest of the time, they declared thinking about something else (Mind Wandering (MW); Face: 38.0 ± 4.3%; Digit: 40.9 ± 4.8%; Figure 1c) or thinking about nothing (Mind Blanking (MB); Face: 12.7 ± 3.0%; Digit: 11.9 ± 2.9%; Figure 1c). These results are well in line with previous findings^34, 35^ and highlight the prevalence of attentional lapses. Attentional lapses were also reflected in participants’ accuracy on the Go trials (Face: 3.1 ± 0.4% of miss, i.e., errors on Go trials; Digit: 1.7 ± 0.2%; Figure 2a), but more notably on the NoGo trials (Face: 35.0 ± 2.5% of false alarms (FA), i.e., errors on NoGo trials; Digit: 32.5 ± 2.7%; Figure 2b).

**Figure 2.**
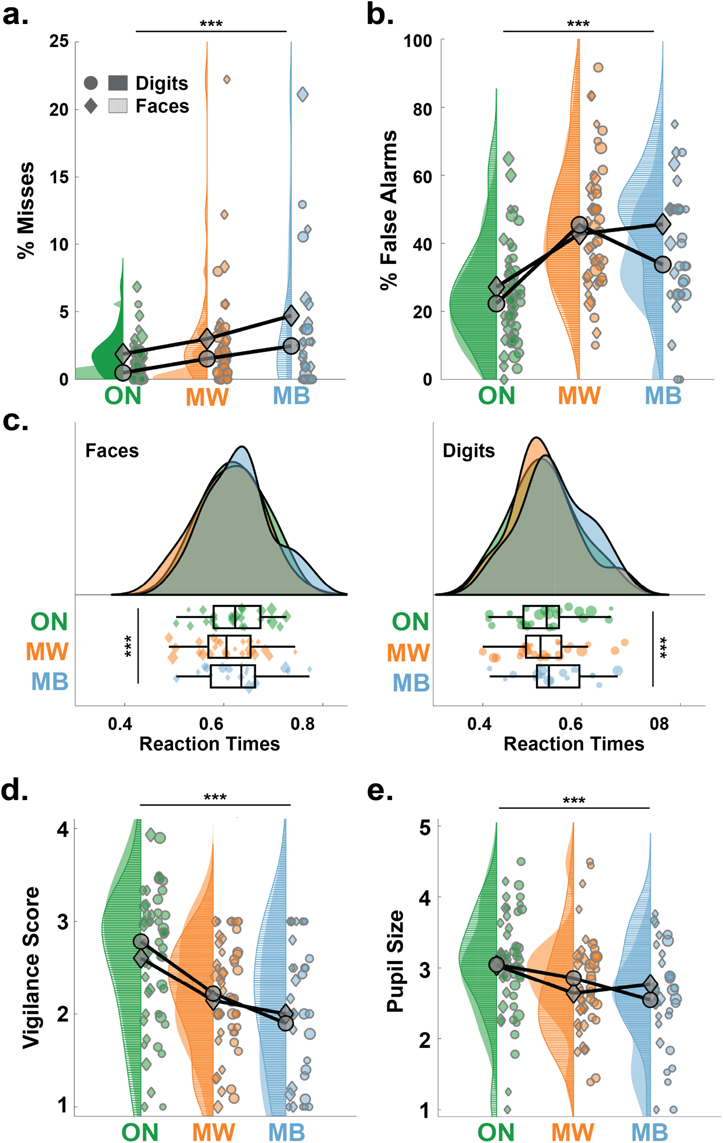
Low arousal is associated with attentional lapses characterized by different behavioural outcomes. Proportion of misses (**a**) and false alarms (**b**) in the 20s preceding task-focused (ON, green), mind-wandering (MW, orange) and mind-blanking (MB, blue) during the tasks with Digits (circles for each individual participant; filled surfaces for smoothed density plot) and Faces (diamonds and surfaces with horizontal stripes). The markers’ size is proportional to the number of reports for each participant (same for **c-e**). Grey diamonds and circles show the average across participants, weighted by the number of reports (same for **c-e**). **c:** Distribution of reaction times (RT) for Go Trials (left: Face; right: Digit) in the 20s preceding ON, MW and MB reports. Box plots show the mean (central bar), the lowest and highest individual data points (end of the whiskers) and the lower and higher quartiles (edges of the box). On top of each plot are shown the smoothed density plot for the different mental states. **d:** Vigilance scores (subjective ratings provided during probes) associated with ON, MW and MB reports. **e:** Discretized pupil size (see Online Methods) in the 20s preceding ON, MW and MB reports. In **a-e**, stars show the level of significance of the effect of mental states (Likelihood Ratio Test, see Online Methods; ***: p<0.005).

Next, we focused on the behavioural patterns preceding subjective reports of attentional lapses. Specifically, we examined participants’ behaviour in the 20s before the onset of the probes that led to MW and MB reports (see Online Methods). To quantify the impact of mental states (i.e., ON, MW or MB) on behaviour, we compared statistical models that either did or did not include mental states as a predictor of behaviour (see Online Methods and Supplementary Table 1). A significantly better fit by the model incorporating mental states, assessed through a Likelihood Ratio Test, was interpreted as evidence for the influence of mental states. To describe the size and direction of the statistical effects, we report the estimates (β) of the contrasts of interest (MW vs ON, MB vs ON and MB vs MW) and their 95% confidence-interval (CI). Accordingly, we found a significant effect of mental states on misses (model comparison: χ^2^(2)=36.0, p=1.5×10^-8^; Figure 2a). Specifically, MW and MB were associated with an increase in misses compared to ON reports (MW vs ON: β=0.011, CI: [0.005, 0.016]; MB vs ON: β=0.023, CI: [0.015, 0.032]) and misses were more frequent for MB compared to MW reports (MB vs MW: β=0.013, CI: [0.005, 0.021]). False alarms were also modulated across mental states (model comparison: χ^2^(2)=115.9, p<10^-16^; see Figure 2b), with an increase for both MW and MB compared to ON (MW vs ON: β=0.21, CI: [0.17, 0.24]; MB vs ON: β=0.17, CI: [0.12, 0.23]), but similar levels of false alarms for MB and MW (MB vs MW: β=-0.028, CI: [-0.084, 0.028]). Finally, mental states were associated with different patterns of reaction times (RT; model comparison: χ^2^(2)=16.3, p=2.9×10^-4^; Figure 2c) with slower reaction times for MB compared to both ON and MW reports (MB vs ON: β=0.019, CI: [0.009, 0.030]; MB vs MW: β=0.022, CI: [0.011, 0.032]; MW vs ON: β=-0.0025, CI: [-0.0096, 0.0045]).

Taken together, these results suggest that MW and MB decrease performance in different ways. MB is characterized with more misses and slower RT than MW and ON, which is consistent with the idea that MB induces sluggish mental states. As to MW, the analysis of misses may ostensibly compatible with an idea of MW as a mild form of MB, however, the fact that RT in MW is faster than in MB is not easy to reconcile with such an idea. Instead, this overall pattern is consistent with an idea that MW induces more impulsive mental states.

The same pattern of results was obtained when normalising misses, false alarms and RT in the MW and MB conditions by the average obtained in the ON condition for each subject (see Supplementary Table 2). We also obtained qualitatively the same results when focusing on a shorter time-window (10s before probe onsets instead of 20s; see Supplementary Table 3).

### Vigilance

Although MW and MB differ according to their phenomenological definition and associated behaviours, both states seem to occur in a similar context of low vigilance (as reported by participants themselves during probes). To address this, we quantified the degrees of correlation between participants’ vigilance ratings and each mental state (comparison between models including or not the information about mental state: χ^2^(2)=144.8, p<10^-16^; Figure 2d and Supplementary Table 1). Participants reported lower vigilance ratings for both MW and MB compared to ON (MW vs ON: β=-0.39, CI: [-0.40, −0.37] and MB vs ON: β=-0.53, CI: [-0.55, −0.50]). Vigilance ratings were even lower for MB compared to MW (MB vs MW: β=-0.13, CI: [-0.24, −0.02]). We then examined a classical objective proxy for vigilance: pupil size^33, 36^. Pupil size prior to probes (Figure 2e, N=25 participants here, see Online Methods) was significantly modulated by mental states (model comparison: χ^2^(2)=18.0, p=1.210^-4^) with MW and MB associated with smaller pupils than ON probes (MW vs ON: β=-0.29, CI: [-0.43, −0.15]; MB vs ON: β=-0.22, CI: [-0.45, −0.003]). Pupil size did not differ between MW and MB (MB vs MW: β=0.065, CI: [-0.16, 0.29]), despite the significant correlation between vigilance ratings and pupil size (model comparison: χ^2^(2)=134.5, p<10^-16^; see Table S1 for details on the models). This implies that these two measures of vigilance can be differentially sensitive to mental states.

### Slow Waves

To further examine the potential mechanistic link between sleep pressure and attentional lapses, we set out to detect a marker of sleep pressure in the EEG signal, in the form of local, sleep-like slow waves. We recently reviewed the rationale behind this approach^11^. In particular, relying on a local marker of sleep allowed us to test whether distinct families of attentional lapses can be coherently explained by the occurrence and spatio-temporal characteristics of slow waves.

Following evidence from ^21, 22, 29, 30^, we operationally defined slow waves as the occurrence of large-amplitude waves within the delta ([1-4] Hz) range. We first detected the occurrence of slow waves in each EEG electrode using an established approach developed in wakefulness and sleep (see ^30, 31, 37^ and Online Methods). Both the temporal profile and topographical distribution of slow waves detected during the tasks (Figure 3a-b) resemble the slow waves observed in NREM sleep^37, 38^. This is not trivial as our detection algorithm did not select this specific temporal profile or topographical distributions.

**Figure 3.**
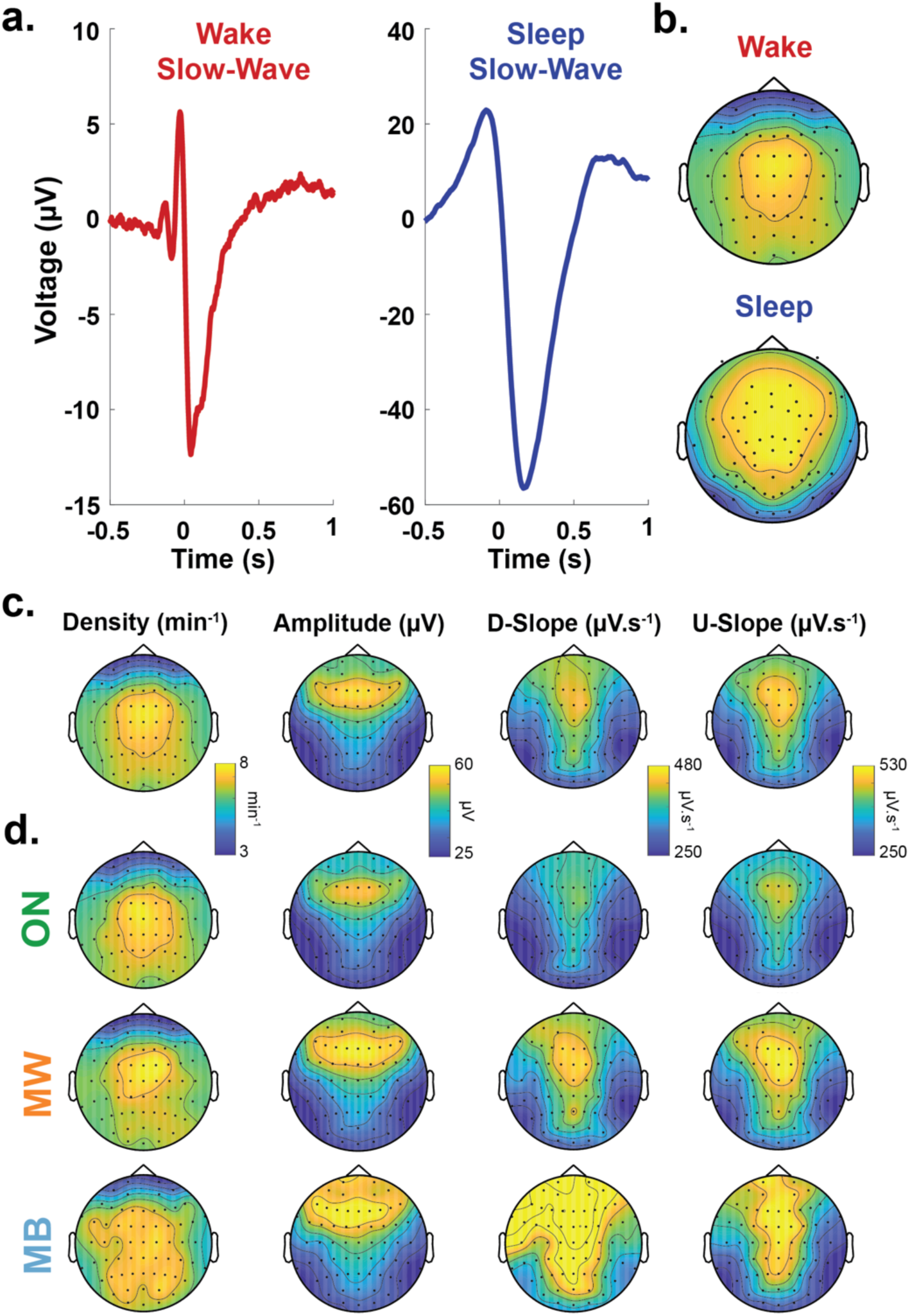
Properties of Slow Waves. **a:** Average waveform of the slow waves detected over electrode Cz during the behavioural tasks (red, left; N=26 participants). The average waveform of slow waves detected during sleep (blue, right), extracted from another dataset (see Supplementary Methods), is shown for comparison. Slow waves were aligned by their start, defined as the first zero-crossing before the negative peak (see Online Methods). **b:** Scalp topographies of the density of slow waves (arbitrary units) detected in wakefulness (top) and sleep (bottom). **c:** Scalp topographies of wake slow-waves properties (1^st^ column: temporal Density; 2^nd^: peak-to-peak Amplitude; 3^rd^: Downward Slope (D-Slope); 4^th^: Upward Slope (U-Slope); see Online Methods) averaged across participants (N=26). **d:** Scalp topographies for slow-waves Density (1^st^ column), Amplitude (2^nd^), Downward Slope (D-Slope, 3^rd^) and Upward Slope (U-Slope, 4^th^) for the different mental state (ON, MW and MB).

#### Relationship between global properties of slow waves and vigilance

Next, we checked whether global properties of the slow waves index participants’ level of vigilance. We correlated first participants’ subjective vigilance ratings with the amount and properties of slow waves. The amount and properties of slow waves were extracted prior to probe onset ([-20, 0]s) and averaged across the entire scalp (all 63 electrodes, see Online Methods). Model comparisons between models with or without slow-wave density, amplitude, downward slope or upward slope indicate a negative correlation between vigilance ratings and slow-waves density (χ^2^(1)=13.1, p=3.9×10^-4^, β=-0.074, CI: [-0.114, −0.034]), amplitude (χ^2^(1)=33.1, p=8.5×10^-9^, β=-0.023, CI: [-0.031, −0.015]), downward slope (χ^2^(1)=82.1, p<10^-16^, β=-2.5×10^-3^, CI: [-3.1×10^-3^, −2.0×10^-3^]), and upward slope (χ^2^(1)=47.3, p<10^-11^, β=-1.7×10^-3^, CI: [-2.2×10^-3^, −1.2×10^-3^]) (see also Figure S1). We then verified that slow-wave properties were negatively correlated with pupil size recorded before each probe (model comparison between models with or without slow-wave density, amplitude, downward slope or upward slope: density: χ^2^(1)=12.3, p=4.6×10^-4^, β=-0.13, CI: [-0.21, −0.059]; amplitude: χ^2^(1)=7.4, p=6.4×10^-3^, β=-0.0088, CI: [-0.015, −0.0025]; downward slope: χ^2^(1)=15.1, p=1.0×10^-4^, β=-6.9×10^-4^, CI: [-1.0×10^-3^, −3.4×10^-4^]; upward slope: χ^2^(1)=17.6, p=2.7×10^-5^, β=-7.6×10^-4^, CI: [-1.1×10^-3^, −4.0×10^-4^]) (see also Figure S2). Finally, we confirmed that the amount of slow waves detected increased with time spent on task (Figure S3), as can be expected when considering the homeostatic regulation of sleep and local sleep^22^.

#### Relationship between local properties of slow waves and mental states at the probe level

Local properties of the slow waves (Figure 3c-d) predicted reports of mental states. Specifically, we examined, for each scalp electrode, whether slow-wave properties (density, amplitude, downward and upward slopes) prior to probes were predictive of the mental state reported by participants. To do so we focused on pairwise contrasts (MW vs ON, MB vs ON and MB vs MW) and performed this analysis for each electrode independently. A cluster permutation approach (p_cluster_<0.05, Bonferroni corrected cluster threshold; see Online Methods) revealed that MW (compared to ON) was predicted by an increase in the number of slow waves and slow wave amplitude over frontal electrodes, and by an increase in slow waves’ downward and upward slope over centro-frontal electrodes (Figure 4a). MB (compared to ON; Figure 4b) was also predicted by an increase in slow wave density over frontal areas as well as by the slow-wave slopes (downward and upward) over centro-parietal electrodes (no significant cluster for slow wave amplitude). Finally, a direct contrast between MB and MW (Figure 4c) indicated that a reduction of slow wave amplitude over frontal electrodes but an increase in their upward slope over parietal electrodes were predictive of MB. No significant clusters were obtained for slow wave density and downward slope.

**Figure 4.**
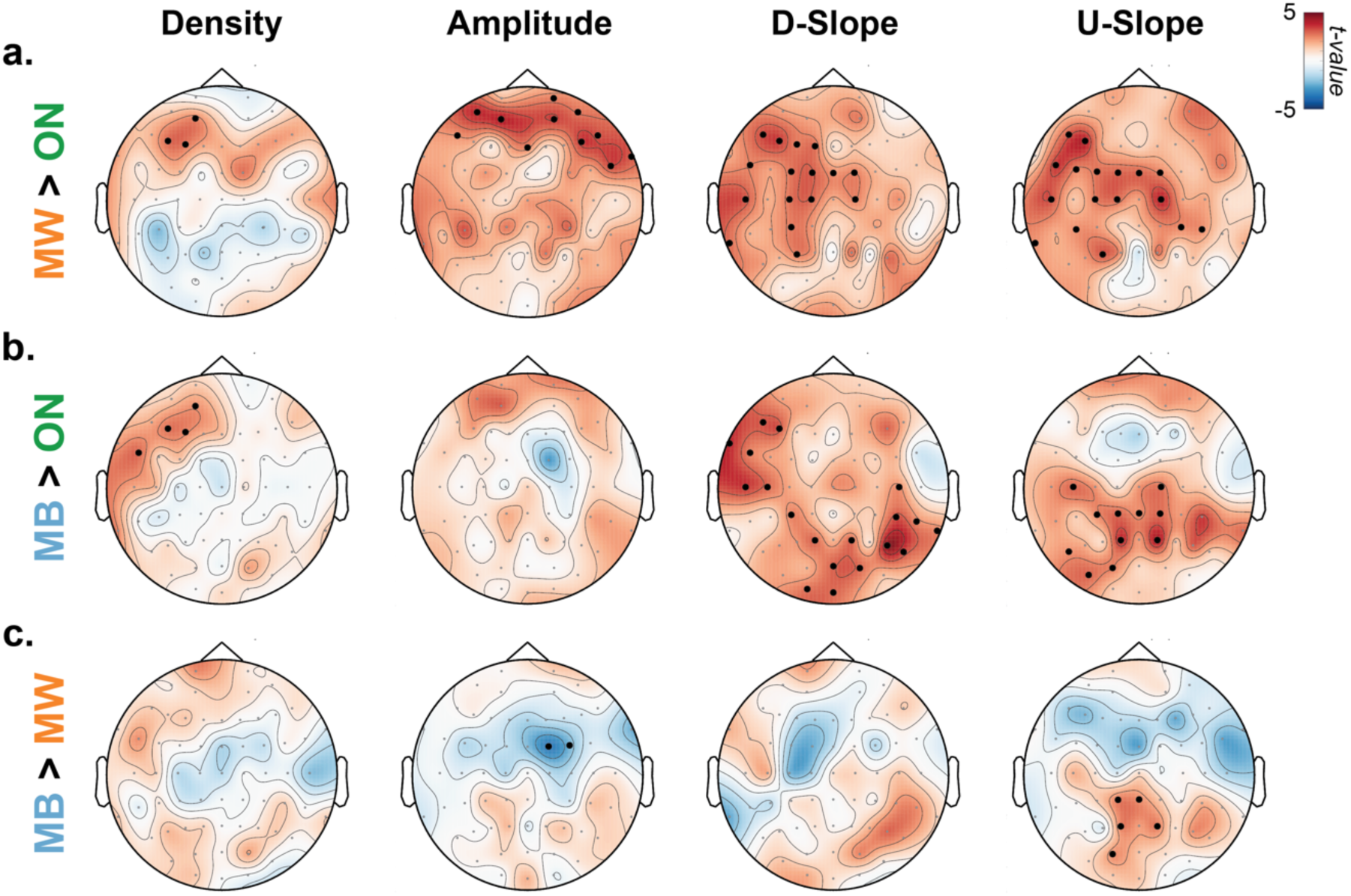
Local properties of slow waves are predictive of mental states. Locally, based on each individual electrode, we performed mixed-effect analyses, following with permutation analysis to quantify the impact of slow wave properties on mental states. 1st column: Density; 2nd: Amplitude; 3rd: Downward Slope (D-slope); 4th: Upward Slope (U-slope). These slow-wave properties were extracted for each electrode and used to compute the t-values (shown in each topographical plot) from t-tests on the following comparisons: (a) MW > ON (b) MB > ON and (c) MB > MW. Black dots denote significant clusters of electrodes (p_cluster_<0.05 corrected for 12 comparisons using a Bonferroni approach, see Online Methods).

To provide more details on the temporal relationship between slow waves and mental states, we replicated this analysis by splitting the 20s window before probes into four windows of 5s (Figure S4-6). The contrasts between MW and ON (Figure S4) as well as MB and ON (Figure S5) show that the properties of slow waves best predict mental states within the 5s before a probe. In terms of topography, when compared to ON state, MW is best predicted by the slow wave properties in frontal electrodes (Figure S4) while MB is best predicted by those in the centro-parietal electrodes (Figure S5).

#### Relationship between local properties of slow waves and behaviour at the trial level

To further understand the association between slow waves and attentional lapses, we examined the influence of slow waves on participants’ behaviour at the single-trial level (for all trials within 20s of a probe onset, independently of mental states). To do so, for each trial and electrode, we marked the presence or absence of slow waves between stimulus onset and offset (see Online Methods) and used this as a (binary) predictor of RT, misses and false alarms (Figure 5). This analysis revealed spatially-specific effects of slow waves on distinct behavioural outcomes. Namely, slow waves in frontal electrodes co-occurred with faster reaction times while slow waves in posterior electrodes co-occurred with slower reaction times (Figure 5a, p_cluster_<0.05, Bonferroni corrected cluster threshold). Likewise, frontal slow waves were associated with more false alarms (a marker of impulsivity, Figure 5b) and posterior slow waves with more misses (a marker of sluggishness, Figure 5c).

**Figure 5.**
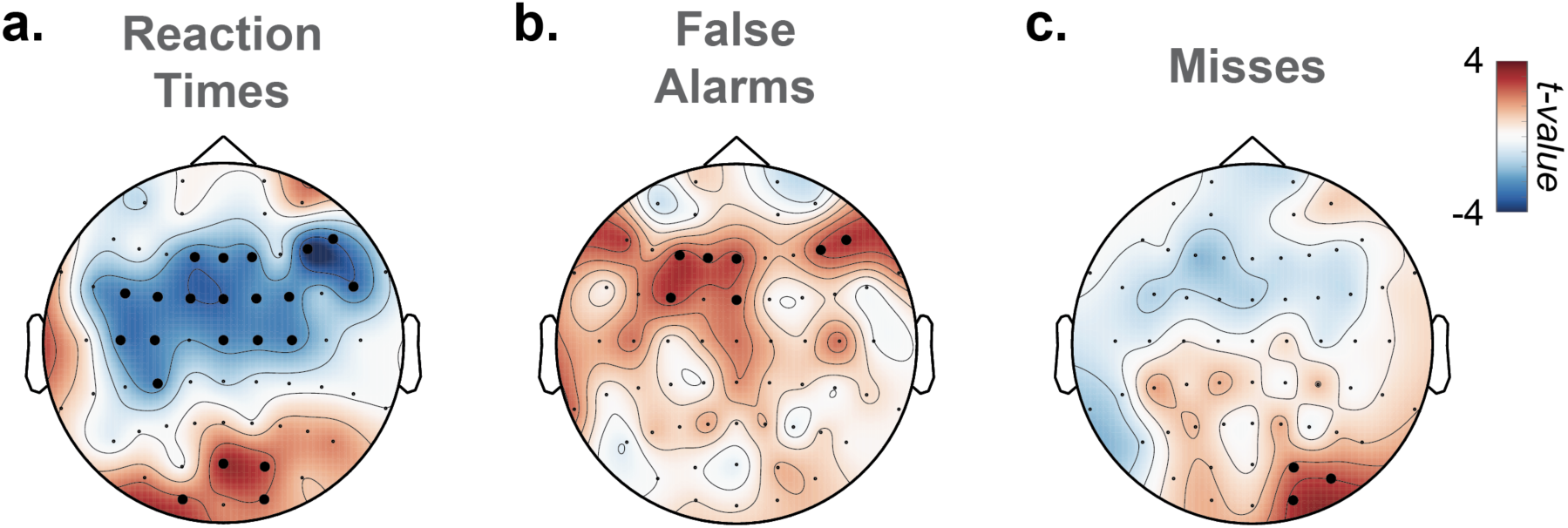
Local occurrences of slow waves are associated with modulations of performance. Mixed-Effects Models were used to quantify the correlation between slow-wave occurrence and reaction times (**a**), false alarms (**b**) and misses (**c**) at the single-trial level. Topographies show the scalp distribution of the associated t-values. Black dots denote significant clusters of electrodes (p_cluster_<0.05 corrected for 3 comparisons using a Bonferroni approach, see Online Methods).

### Decision modelling

Finally, we implemented an influential model of two-alternative forced-choice (2AFC) decision making: the diffusion decision model (DDM)^39^. The DDM decomposes full reaction-time distributions and choice proportions into latent cognitive processes that are thought to underlie participants’ decisions in 2AFC tasks (see Online and Supplementary Methods and Figure S7). These include the time it takes for participants to start computing their response (Non-Decision Time or NDT [*t*]), the speed at which participants accumulate evidence for the two responses (drift rates for Go [*v_Go_*] and NoGo [*v_NoGo_*] responses), the amount of evidence needed to reach a decision (decision threshold [*a*]), the participants’ initial bias for one of the two responses (decision bias [*z*]) and the bias for the accumulation of evidence for one of the two responses (drift bias, [*v_Bias_*]). A hierarchical Bayesian approach was used to fit the DDM to the reaction times obtained in the Go/NoGo tests^40^ so that each parameter (*v_Go_*, *v_NoGo_*, *a*, *z*, *t* and *v_Bias_*) was free to vary by participant, task and mental states (ON, MW and MB) or slow-wave occurrence (present vs. absent). Simulations confirmed this hierarchical DDM can successfully predict the observed data (Figure S8).

Considering the trials that were within 20 seconds from the onset of the probes, we first estimated these different parameters of the DDM for each mental state (Figure S9). This analysis shows differences between MW and MB reports in terms of decision bias and threshold (Figure S9). The lower threshold and higher decision bias observed for MW (compared to MB) are concordant with the idea that MW is associated with behavioural impulsivity.

We then used this modelling approach to examine how the occurrences of slow waves impact the different cognitive processes leading to participants’ responses, with a particular focus being the test of our core hypothesis: the presence of slow waves in frontal areas leads to impulsivity by disrupting the cognitive mechanisms underlying executive control, while the presence of slow waves in posterior areas leads to sluggishness by slowing down the integration of sensory inputs.

We report here the differences in the parameters’ estimates in the presence or absence of slow waves detected for each electrode (Figure 6). The associated scalp topographies indicate both global and local (electrode-specific) effects of slow waves on DDM parameters. As global effects (Figure 6d-f), we found first that the presence of slow waves was associated with a reduction in decision threshold (*a*; Figure 6d), consistent with the idea that slow waves facilitate impulsive responses. Second, slow waves were also associated with longer non-decision times (*t*; Figure 6e), suggesting that slow waves can slow down neural processes underlying stimulus encoding and/or motor preparation. Finally, the presence of slow waves was correlated with an increase in the starting point of the decision process (prior bias *z*; Figure 6f), implying shifts in the decision process towards Go responses.

**Figure 6.**
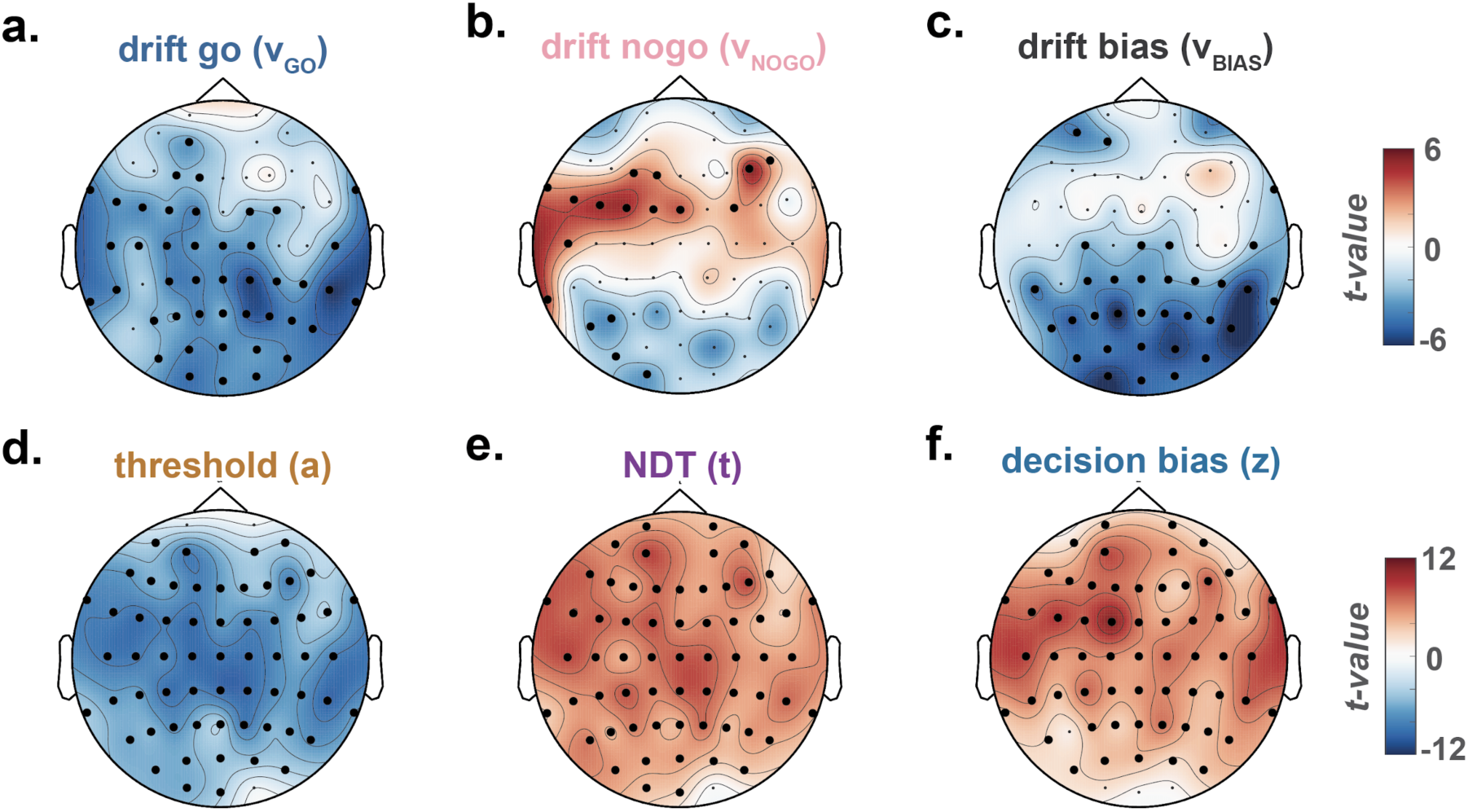
Global and local effects of the occurrence of slow waves on sub-components of decision-making. Reaction Times in the Go/NoGo tasks were modelled according to a Hierarchical Drift Diffusion Model (see Online Methods). a-f: Topographical maps of the effect of slow waves (i.e. whether or not a slow wave was detected for each trial and for a specific electrode) on the parameters of decision-making: drift Go [v_Go_] (a), drift NoGo [v_NoGo_] (b), drift bias [v_Bias_] (c), threshold [a] (d), Non-decision time or NDT [t] (e), decision bias [z] (f). The effect of slow wave occurrence was estimated with LMEs (see Online Methods) and topographies show the scalp distribution of the associated t-values. Black dots denote significant clusters of electrodes (pcluster<0.05, Bonferroni-corrected, see Online Methods).

As local electrode-specific effects, we observed again contrasting results between posterior and frontal slow waves (Figure 6a-c). Slow waves within posterior electrodes were associated with a reduction of *v_Go_* and *v_Bias_*, meaning that evidence accumulation was slower for Go decisions and the drift bias of the decision process for Go responses was reduced (Figure 6a,c). Conversely, slow waves within frontal electrodes correlated with a reduction of *v_NoGo_*, indicating that evidence accumulation was slower for NoGo Decisions (Figure 6b). These modelling results provide a potential explanation on how local slow wave properties relate to mental states (Figure 4) and single-trial task performance (Figure 5). In other words, posterior slow waves were associated with sluggish responses and increased misses while frontal slow waves were associated with faster, impulsive responses and more false alarms. Indeed, slower evidence accumulation in favour of Go responses would lead to slower reaction times or even misses, whereas slower evidence accumulation for NoGo responses would lead to faster responses and possibly false alarms.

These results suggest that slow waves could represent both a general index of fatigue as well as a mechanism underlying specific behavioural and subjective consequences of sleepiness. In other words, regardless of where it happens, slow waves can reflect the global “sleepiness” of an individual. Meanwhile, at a finer grain, depending on where they occur, slow waves can disrupt specific cognitive processes carried out by the affected brain regions. Taken together, we interpret this as strong evidence in support of the idea that slow waves are a compelling physiological phenomenon, which precedes and co-occurs with subjective and behavioural aspects of attentional lapses. In addition, our results indicate that spatio-temporal properties of slow waves can specifically explain distinctive components of behavioural and mental consequences of attentional lapses in a unified and quantitative manner.

## Discussion

According to both in-lab and real-life studies, humans spend up to half of their waking lives not paying attention to their environment or any task at hand^34, 35^. However, the ubiquitousness of these attentional lapses remains unexplained. Part of the difficulty in identifying their neural correlates could be due to their intrinsic diversity^7, 8^. We embraced this challenge to clarify the neural mechanisms underlying attentional lapses by linking three different levels of explanation: behaviour, phenomenology and physiology. To grasp the phenomenological diversity of attentional lapses, we distinguish between MW and MB, where MW is operationally defined as a shift in attention away from environmental demands and toward spontaneous task-unrelated thoughts, and MB as the absence of thoughts and an empty mind. We propose that these two types of attentional lapses can be explained by the occurrence of sleep-like low-frequency high-amplitude waves, where regional differences in the occurrence of slow waves predicts the type of attentional lapse as well as its behavioural profile. These slow waves have been previously linked to the transition toward sleep when they are visible across the scalp^38^ or to the concept of local sleep when they occur in a subset of electrodes^11, 22, 25, 29, 30^ and in the context of an awake individual.

The concept of local sleep builds upon a recent questioning of the classical view of sleep as an all-or-nothing phenomenon^22–24, 41, 42^. Although sleep is orchestrated at the scale of the entire brain, some of the key neural mechanisms underlying sleep are implemented and regulated at the level of local cortical circuits^24, 25^. Consequently, when the pressure for sleep increases, an awake brain can start displaying local sleep-like patterns of activity such as delta or theta waves^21, 22, 29, 43^. These bouts of local sleep-like activity, in the form of large-amplitude slow waves, are both time-dependent (i.e. increase with time spent awake) and use-dependent (i.e. depend on the level of activation of a given brain region)^29, 30^. The occurrence of slow waves has been linked to perturbations of information processing and task-related errors or attentional lapses in animal and human intracranial data^21, 22^. A similar relationship between local sleep and behavioural errors has been evidenced when detecting slow waves in non-invasive scalp recordings of sleep-deprived^29, 30^ or well-rested^31^ individuals.

Importantly, our study proposes a novel link between sleep-like slow waves in normal wakefulness and both behaviour and subjective experience. Specifically, we showed that, within the same individual, slow waves temporally precede different types of attentional lapses. At the behavioural level, we observed that sluggish responses (slow responses and misses) were associated with an increase in slow waves over posterior electrodes (Figure 5a,c). Conversely, impulsive responses (hasty responses and false alarms) were associated with an increase in slow waves over frontal electrodes (Figure 5a-b). At the phenomenological level, compared to task-focused (ON) reports, mind wandering (MW) was preceded by an increase in slow-wave density, amplitude, upward and downward slopes over frontal electrodes (Figure 4a). Compared to ON reports, mind blanking (MB) was also preceded by an increase in slow-wave density over frontal electrodes, but we additionally observed an increase in slow-wave upward and downward slopes over posterior electrodes (Figure 4b). A direct comparison between MW and MB reports shows that MB reports were associated with slow waves with steeper upward slopes over posterior electrodes and smaller amplitude over frontal electrodes (Figure 4c). Finally, focusing on shorter time windows (Figure S4-6), we showed that slow waves best predict mental states (MW or MB vs ON) in the last few seconds before a probe, in line with previous findings^44, 45^.

These results imply a spatial and temporal relationship between markers of local sleep and behavioural errors: local sleep events, as measured through the presence of slow waves, occurring at the right time (i.e. during stimulus presentation) and in the right place (i.e. in the brain regions involved in the task) are predictive of distinct behavioural and phenomenological aspects of attentional lapses^11^.

Importantly, our results are largely consistent with previous findings on the neural correlates of attentional lapses. Most of these studies focused on mind wandering, although often defined as any mental state that is not on task (i.e. attentional lapses, which include MW and MB as defined here). Early fMRI studies showed that mind wandering in this sense was associated with the activation of the Default Mode Network (DMN)^46, 47^. Relevant to our local sleep interpretation, the DMN is also suggested to be involved in dream generation during sleep, consistent with the idea that the DMN supports a broad array of experiences that are decoupled from the environment^48^. Furthermore, lesions within the DMN are associated with a decrease in both mind wandering^49^ and dreaming^50^. However, recent findings suggest a complex relationship between the DMN and spontaneous experiences. Unlike what was initially hypothesized, the DMN is now considered to be involved in both task-unrelated and task-related processes^44, 51^, depending for example on environmental demands or the vividness of individuals’ experiences^52, 53^. Based on our results, we speculate that local sleep plays a key role in the complex relationship between the DMN and spontaneous experiences. Previous studies have indeed shown that a state of low alertness could induce the phasic activation of the DMN^54^. We speculate that slow waves occurring within the DMN could lead to episodes of mind wandering by disconnecting individuals from their environment^55^. Indeed, slow waves during sleep have been proposed to be responsible for disconnecting sleepers from their environment^27, 56^, as they are accompanied by a phenomenon of neuronal silencing that can disrupt the processing of external inputs. If slow waves trigger specific occurrences of mind wandering, other mechanisms could be responsible for the stability of these episodes^57^. Further investigations, including source localization or simultaneous recording of EEG and fMRI^58, 59^, promise a deeper understanding of the mechanisms underlying these attentional lapses.

Our work might also help reconcile previous findings on the EEG correlates of mind wandering. A seminal study reported a reduction of alpha and/or beta oscillations during mind wandering,^60^ whereas others have reported an increase^61–63^. However, the relation between alpha and sleep is complex: alpha power is low both when participants are fully alert and, on the contrary, when approaching sleep onset^64, 65^. Perhaps, the divergent results obtained in previous studies regarding mind wandering and alpha oscillations could be explained by different baseline levels of alpha oscillations in these studies. Likewise, past pupilometry results on attentional lapses have been similarly inconclusive. While most studies found a dampening of stimulus-locked increases in pupil size during mind wandering (e.g. ^47, 66, 67^), results diverge for baseline pupil size, which has been reported as either increasing (e.g. ^68, 69^) or decreasing (e.g. ^47, 70, 71^). When distinguished from mind wandering, mind blanking has also been associated with reduced pupil size compared to task-focused states^10, 66^. Our results largely align with the latter results, with both mind wandering and mind blanking being characterized by a decrease in pre-probe pupil size (Figure 2e), which goes together with low vigilance ratings (Figure 2d) and an increase in slow wave occurrence (Figure 3-4). In addition, the complex relations between pupil size and mind wandering may be partly explained by the fact that pupil size does not index only arousal but also correlates with motivation^72^ and cognitive load^73^, which also correlate with mind wandering^2^. In contrast, the sleep-like nature of wake slow waves would make them an unambiguous marker of sleepiness. Furthermore, slow waves are detected at the electrode level and can therefore indicate how brain regions respond differently to sleep pressure.

Other than mind wandering and blanking, we foresee that the spatio-temporal properties of slow waves might predict other types of spontaneous experiences. For example, previous research focusing on the hypnagogic period at sleep onset has shown that slow-wave like activities are predictive of the occurrence of spontaneous imagery^74^, the intensity of thoughts^58^ and can discriminate between different contents of spontaneous experiences^75^. Likewise, during sleep, it has been reported that local modulation of slow wave power is predictive of the occurrence and contents of dreams^76^. Local sleep, therefore, might be related to the type and content of spontaneous experiences not only during wakeful states, as we showed here, but also during sleep-wake transitions and sleep.

Our results also speak to the broader question of how different brain regions participate in shaping the stream of consciousness. Slow waves, considered as a perturbation of the normal wake activity of local cortical networks, could reflect the functions of these networks under normal situations. For example, we observed that slow waves in frontal regions were associated with false alarms (Figure 5b), which aligns well with the role of frontal cortices in executive functions and response inhibition^77^. Conversely, slow waves in the back of the brain were associated with misses (Fig 5c), which is consistent with the involvement of parietal cortices in sensorimotor integration^78^. At the phenomenological level, frontal slow waves were associated with mind wandering, whereas posterior slow waves were associated with mind blanking. Thus our results could speak to the debate on the respective involvement of frontal and posterior cortices in supporting different conscious states^79, 80^. Our results suggest that a perturbation of frontal cortices leads to unconstrained thoughts (mind wandering) rather than the loss of awareness, but that awareness decreases when posterior regions momentarily go offline, a pattern similar to the neural correlates of dreaming during sleep or spontaneous thoughts during wakeful rest^58, 76^. However, frontal and posterior slow waves do not only differ by their location but also in terms of spatial expanse: slow waves in frontal electrodes appears more focal, whereas slow waves in posterior electrodes is more widespread (Figure S10). Thus, the loss of awareness during mind blanking could reflect a tendency for posterior slow waves to involve a broader fronto-parietal network. This is in line with theories attributing an essential role of fronto-parietal connections in the emergence of consciousness^81–83^.

We speculate that the slow waves we report here are generated by similar neural mechanisms as slow waves in sleep. Our speculation rests on the profile of slow waves characterized in (i) time (Figure 3a) and (ii) space (Figure 3b,c), and the relationships between slow wave properties and (iii) time spent on task (Figure S3), (iv) subjective vigilance and (v) pupil size. While these five lines of evidence are correlational, we note that together, they imply a high degree of similarity between slow waves in waking and sleep. This, in turn, allows us to interpret wake slow waves as local sleep. Importantly, as our study builds on convergent lines of indirect evidence to argue that the slow waves we measured in waking participants are sleep-related, this interpretation is tentative and we do not wish to suggest that the presence of slow-frequency oscillations in wake EEG is by itself sufficient to establish a relation to sleep. Here, for example, we paired these observations with objective (pupil size) and subjective (vigilance ratings) markers of fatigue and checked that slow waves increased with the time spent on task. Future studies could use direct evidence from intracranial recordings or sleep deprivation to more solidly establish this interpretation. Intracranial recordings in particular can confirm whether wake slow waves are accompanied by episodes of neuronal silencing, as for sleep slow waves, and would help the identification of the exact neural mechanisms that generate the slow waves we report here.

As our task involved the continuous presentation of visual stimuli, we could not fully disentangle the occurrence of slow waves and task-related events. Yet, we showed that slow waves differ from typical responses evoked by stimuli or participants’ responses (Figure S11). Future studies could include task-free resting state sessions to compare the occurrence and properties of slow waves occurring during a task and without a task. Relatedly, further investigation could help determine to which extent our findings generalize to everyday situations (e.g., driving, attending a lecture, reading, etc).

It is worth noting that the tasks used here (SARTs) are rather undemanding and could favour sleepiness and local sleep compared to more difficult and engaging experimental paradigms. Generalising our findings to different experimental contexts or more naturalistic settings would need further experimental validation^11^. From our findings, we predict that slow waves would predict the occurrence of attentional lapses and mind wandering in situations where participants feel rather sleepy and to a lesser extent or not at all in situations where participants are well awake, motivated and highly engaged. We also speculate that slow waves are not the only mechanism that underlies sensory decoupling in wakefulness and sleep^56^. Other factors, such as the neuromodulation of brain activity, may be critical in clarifying the total set of mechanisms underlying mind wandering^9, 84^ and blanking, and more broadly attentional lapses.

In conclusion, we show here that attentional lapses occurring in the context of an undemanding task are accompanied by sleep-like slow waves, even when participants are well-rested. Furthermore, the location of slow waves is predictive of certain behavioural and phenomenological properties of these lapses. Thus, we propose that these slow waves reflect local intrusions of sleep within waking and constitute a mechanistic and proximate cause to explain attentional lapses. Identifying a proximate mechanism of attentional lapses could inspire novel applications leveraging brain-machine interfaces in educational or professional settings.

## Online Methods

### Participants

Thirty-two (N=32) healthy adults were recruited and participated in this study. Six individuals were not included in our analyses because of technical issues during recordings or an abnormal quality of physiological recordings assessed through a post-hoc visual inspection of the data. The remaining 26 participants (age: 29.8 ± 4.1 years, mean ± standard-deviation; 10 females) were included in all analyses except for one individual for whom we do not have eye-tracking data.

### Experimental Design and Stimuli

Participants were seated in a dimly lit room with their chin resting on a support at approximately 60cm from a computer screen. All task instructions and stimuli were displayed and button responses were collected via the Psychtoolbox extension^85^ for Matlab (Mathworks, Natick, MA, USA).

The experimental design consisted of two modified Sustained Attention to Response Tasks (SARTs)^32^ in which participants were instructed to pay attention to a series of pictures of human faces in the Face SART blocks or digits in the Digit SART blocks. The order of Face or Digit blocks was pseudo-randomized for each participant. Each block lasted approximately 12 to 15 minutes. Participants were allowed to rest between blocks. Participants performed 3 Face SART blocks and 3 Digit SART blocks for a total duration of 103 ± 19.7 minutes (mean ± standard-deviation) from beginning to end. Each type of the Face and Digit SART was preceded by a brief training session (27 trials) on each SART. During this SART training session, feedback on the proportion of correct trials and average reaction times was provided to participants. Participants were encouraged to prioritize accuracy over speed.

Face and digit stimuli were presented continuously, each stimulus appearing for a duration of 750 to 1250ms (random uniform jitter). Face stimuli were extracted from the Radboud Face Database^86^ and consisted of 8 faces (4 females) with a neutral facial expression and one smiling female face. Digits from 1 to 9 were displayed with a fixed font size. For the Face SART, participants were instructed to press a button for all neutral faces (Go trials) but to avoid pressing the response button for the smiling face (NoGo trials). The order of faces was pseudo-randomized throughout the entire task (i.e. we permuted the presentation order every 9 stimuli and we did not present twice the same stimuli in a row). For the Digit SART, participants were instructed to press a button for all digits except the digit “3” (NoGo trials), with the order of the digits pseudo-randomized as well.

During the SART, we stopped the presentation of stimuli at random times (every 30 to 70s, random uniform jitter) with a sound and the word “STOP” displayed on the screen. These interruptions allowed us to probe the mental state of the participants with a series of 8 questions (including 1 conditional question; see Supplementary Methods). In particular, we instructed participants to report their attentional focus “just before the interruption”. Participants had to select one of the four following options: (1) “task-focused” (i.e. focusing on the task, ON), (2) “off-task” (i.e. focusing on something other than the task, which we define here as mind wandering MW), (3) “mind blanking” (i.e. focusing on nothing), (4) “don’t remember”. As the 4th option accounted for only 1.1% of all probes (i.e. less than 1 probe per participant on average) and since previous studies do not always distinguish between these options (e.g. ^87^), we collapsed the 3^rd^ and 4^th^ options as mind-blanking (MB) in all analyses. We also instructed participants to rate their level of vigilance, reflecting “over the past few trials”, with a 4-point scale (Figure 2d; from 1=“Extremely Sleepy” to 4=“Extremely Alert”). Each of the 12-15 min SART blocks included 10 interruptions (in total, 30 interruptions for each SART task and 60 interruptions per participant). Participants were informed of the presence of interruptions and the nature of each question before starting the experiment. The mental state categories (ON, MW and MB) were also explained to participants orally and in writing.

### Physiological Recordings and Preprocessing

High-density scalp electroencephalography (EEG) was recorded using an EasyCap (63 active electrodes) connected to a BrainAmp system (Brain Products GmbH). A ground electrode was placed frontally (Fpz in the 10-20 system). Electrodes were referenced online to a frontal electrode (AFz). Additional electrodes were placed above and below the left and right canthi respectively to record ocular movements (electrooculogram, EOG). Two electrodes were placed over the deltoid muscles to record electrocardiographic (ECG) activity. EEG, EOG and ECG were sampled at 500Hz. Eye-movements and pupil size on one eye were recorded with an EyeLink 1000 system (SR Research) with a sampling frequency of 1000Hz. The eye-tracker was calibrated at the start of each recording using the EyeLink acquisition software.

The EEG signal was analysed in Matlab with a combination of the SPM12, EEGlab^88^ and Fieldtrip^89^ toolboxes. The raw EEG signal was first high-pass filtered above 0.1 Hz using a two-pass 5^th^-order Butterworth filter. A notch filter was then applied (stop-band: [45, 55] Hz, 4^th^-order Butterworth filter) to remove line noise. Electrodes that were visually identified as noisy throughout the recording were removed and interpolated using neighbouring electrodes. Finally, the continuous EEG data was segmented according to probe onsets on a 64s window ([-32, 32] s relative to the probe onset); the average voltage over the entire window (64s) was then removed for each electrode and probe.

Pupil size was analysed with custom functions in Matlab and corrected for the occurrence of blinks (see ^33^ and Supplementary Methods). Pupil size was averaged over the stimulus presentation window for each trial (window length: 0.75 to 1.25s). Pupil size values in Figure 2e were computed by averaging the pupil size in all trials within 20s preceding the probe onset and then by discretizing them into 5 bins across all probes for each participant and task, to normalize pupil size across participants^33^.

### Behavioural Analyses

Go trials were considered incorrect (Miss) if no response was recorded between stimulus onset and the next stimulus onset. Conversely, NoGo trials were considered incorrect (false alarm) if a response was recorded between stimulus onset and the next stimulus onset. Reaction times were computed from the onset of the stimulus presentation. Trials with reaction times shorter than 300ms were excluded from all analyses (so not considered correct or incorrect). In all subsequent analyses, we focused only on trials within 20s from probe onset. The choice of a 20s-window allowed the inclusion of 2 NoGo trials for each probe, while focusing on trials that are relatively close to the probe onset and subsequent subjective reports. Since previous studies sometimes examined shorter pre-probe time windows (e.g. ^44^), we also analysed the data with shorter time windows (Table S3 and Figure S4-6).

### Detection of Slow Waves

The detection of sleep-like slow waves in waking was based on previous algorithms devised to automatically detect slow waves during NREM sleep^30, 37^. First, the preprocessed EEG signal was re-referenced to the average of the left and right mastoid electrodes (TP9 and TP10) to match the established guidelines for sleep recordings^90^. Then, the signal was down-sampled to 128Hz and band-pass filtered within the delta band. A type-2 Chebyshev filter was used to reach an attenuation of at least 25 dB in the stop-band ([0.1, 15] Hz) but less than 3 dB in the pass-band ([1, 10] Hz). All waves were detected by locating the negative peaks in the filtered signal. For each wave, the following parameters were extracted: start and end point (defined as zero-crossing respectively prior the negative peak of the wave and following its positive peak), negative peak amplitude and position in time, positive peak amplitude and position in time, peak-to-peak amplitude, downward (from start to negative peak) and upward (from negative to positive peak) slopes.

Slow waves in sleep typically have a larger negative peak compared to their positive peak (Figure 3a) and are predominantly observed over fronto-central channels^37, 38^ (Figure 3b). This contrasts with artefacts in the EEG signal caused by blinks, which typically have a large positive component and are more frontally distributed. To reduce the false detection of these artefacts as candidate slow waves, we excluded waves with a positive peak above 75µV. We also excluded waves within 1s of large-amplitude events (>150µV of absolute amplitude). Finally, we discarded all waves that were shorter than 143ms in duration (corresponding to a frequency > 7Hz). We then selected the waves with the highest absolute peak-to-peak amplitude (top 10% computed for each electrode independently) as local sleep slow waves. This 10% threshold was selected based on the visual examination of the distribution of the amplitudes at the subject level (see Figure S12 for the corresponding topography). The mean ± SEM of the threshold voltage across subjects was 30.7 ± 1.5 µV. Finally, we compared slow waves with neural activity related to task events (stimulus onsets and motor responses). To do so, we computed the event related potentials (ERPs) by averaging, across participants, the EEG signal band passed around 0.1 and 30Hz and referenced to the mastoids. These ERPs were locked either on slow-waves’ start, stimulus onset (for face and digit stimuli separately) or motor responses. The corresponding temporal profiles are shown in Figure S11a-c for electrode Cz.

### Hierarchical Drift Diffusion Modelling

Hierarchical Bayesian Drift Diffusion Modelling (HDDM) was used to extend our analysis beyond simple behavioural metrics and examine the impact of mental states (Figure S9) and slow waves (Figure 6) on the sub-processes of decision making. The DDM is a sequential-sampling model of 2AFC decision making that can be considered an extension of signal detection theory into the time-domain, accounting for full reaction time distributions as well as choice behaviour^39^. The HDDM package^40^ in Python 2.8 was used to fit the drift-diffusion model to the SART data. DDM parameters were estimated using a hierarchical Bayesian method that uses Markov-chain Monte Carlo (MCMC) sampling to generate full posterior distributions of model parameters. The following DDM parameters were estimated: the drift rate for Go trials (*v_Go_*), the drift rate for NoGo trials (*v_NoGo_*), the decision threshold (*a*), the decision bias (*z*) and the non-decision time (*t*). Drift bias (*v_Bias_*) was computed by taking the difference between the absolute values of *v_Go_* (positive) and *v_NoGo_* (negative), where greater values indicate stronger *v_Go_* drift bias (Figure S7). To examine whether the model could reproduce key patterns in the SART data, posterior predictive checks were undertaken by simulating 100 datasets from the joint posteriors of model parameters and comparing these to the observed data^91^ (Figure S8).

To estimate HDDM parameters 8,000 samples from the posterior were generated with MCMC methods and the initial 2,000 were discarded as burn-in to reduce autocorrelation. HDDM models were first fit on the trials preceding subjective reports to examine the influence of mental states on decision parameters (Figure S9). HDDM parameters were free to vary by state and task. From the estimated models, we extracted the subject-level point-estimates of parameters as the mean of each individual’s posterior distribution for a given task (Face and Digit) and mental state (ON, MW or MB). We then fitted HDDM models so as to examine the influence of slow waves (event present vs. absent; Figure 6). To do so we considered each EEG electrode separately. For a given electrode, a trial was flagged as being associated with slow waves if the onset of a slow wave was detected for this electrode during stimulus presentation (i.e. between stimulus onset and offset). Parameters were also free to vary by task (Digit vs. Face). When examining the impact of mental states or slow waves on HDDM parameters, we included trials within 20s of a probe onset (allowing to include 2 NoGo trials for each probe). From the estimated models, we extracted the subject-level point-estimates of parameters as the mean of each individual’s posterior distribution for a given task (Face and Digit) and slow waves (present or absent). Statistical comparisons were performed on the subject-level point-estimates.

### Statistical Analyses

Statistics were performed using Linear Mixed-Effects modelling (LMEs). In all models, subject identity was coded as a categorical random effect. The task type (Digit or Face SART) was used as a categorical fixed effect in all analyses. Several fixed effects were independently tested in our different analyses: mental state (categorical variable: ON, MW and MB; Figure 2-4) or slow waves (binary variable: present/absent, Figure 5). LMEs were run to predict different variables of interest: behavioural variables (misses, false alarms, reaction times) or physiological variables (pupil size, presence or properties of slow waves). We also used LMEs to estimate the effect of mental states or slow waves on the point-estimates derived from the HDDM models (Figure S9 and Figure 6). Model comparisons were performed using a Likelihood Ratio Test to estimate the influence of multi-level categorical variables such as mental states. In practice, we compared a model including mental state (along possibly other random and fixed effects) with a model not including mental state as a predictor. All models and model comparisons are described in Supplementary Table 1. In the Results section, we report the Likelihood Ratio Test as χ^2^(df), where χ^2^ is the Likelihood Ratio Test statistic and df the degrees of freedom^92, 93^. When several model comparisons were performed for the same analysis using the Likelihood Ratio Test, a Bonferroni correction was applied to the statistical threshold. To indicate the magnitude and direction of the effects, we report the estimates (β) and confidence interval (CI) for the contrasts of interest (MW vs. ON, MB vs ON, MB vs MW). All models performed are described in Supplementary Table 1. For topographical maps, clusters were identified using a cluster-permutation approach. In practice, for each electrode, we extracted the t-values and p-values for the effects of interest. Clusters of neighbouring electrodes were defined as electrodes with p-values below 0.025 (cluster alpha). Once the clusters were defined, a comparison was performed with permuted datasets and we used a Monte-Carlo p-value threshold of 0.05 to identify the significant clusters (p_cluster_, see Supplementary Methods for details). This threshold was corrected for multiple comparisons when several non-independent cluster permutations were performed (e.g. Figure 4-6, S2-4) in order to keep the type-1 error rate constant across analyses.

## Acknowledgments

TA and NT were supported by Australian Research Council Discovery Projects (DP180104128 and DP180100396) and National Health Medical Research Council Ideas Projects (APP1183280). TA was supported by a Long-Term Fellowship from the Human Frontier Science Program (LT000362/2018-L). JW was supported by Australian Research Council Discovery Early Career Researcher Awards (DE170101254). We thank Devangna Tangri for her help in data collection and Giulio Bernardi for sharing his slow-wave detection algorithm. We also thank our reviewers, Dr Tristan Bekeinschtein, Dr Jonathan Smallwood and a third anonymous reviewer for their constructive comments and criticisms.

## Author Contributions

Design: TA, TM, JW & NT. Data collection: TA & TM. Analyses: TA & AB. Manuscript: TA, AB, JW & NT.

## Data Availability

All raw data and code generated for this study are publicly available:

Raw data: https://osf.io/ey3ca/?view_only=680c39e7065649c3b783a4efec0a1a94

Code used for analyses: https://github.com/andrillon/wanderIM

## Supplementary Information

### Supplementary Methods

#### Participants

Prior to their participation in the protocol, all 26 participants but one filled in online surveys on Qualtrics (N=25). They reported normal levels of sleepiness (Epworth Sleepiness Scale: 14.6 ± 4.7; mean ± standard-deviation) and mind wandering (Mind Wandering Questionnaire^1^: 3.6 ± 0.91) in their everyday lives. Participants were not instructed to follow a particular schedule before the experiment (in particular, our protocol did not involve a sleep restriction procedure). The timing of the experiment was determined by participants’ availability and included both morning and afternoon sessions.

#### Experimental Design and Stimuli

Face stimuli were divided in two parts vertically (half-left and half-right faces) which were flickered on the screen at different frequencies (12 and 15 Hz, counterbalanced across participants). Similarly, the digits were inserted in a Kanizsa illusory square (Fig. 1a) whose right and left parts also flickered at different frequencies (12 and 15 Hz). This flickering was introduced to elicit Steady State Visual Evoked Potentials (SSVEPs) in the EEG signal. The detailed analysis of this aspect of our dataset will be reported elsewhere. As the flickering occurred at a high rate, participants did not report a negative effect on their ability to perform the SART.

#### Experience Sampling

Following task interruptions (probes), participants were asked to answer a series of 8 questions in the following fixed order: (1) ‘Were you looking at the screen?’ (response: yes / no); (2) ‘Where was your attention focus?’ (response: on-task / off-task / blank / don’t remember); (3) ‘What distracted your attention from the task?’ (response: Something in the room / personal / about the task); (4) ‘How aware were you of your focus?’ (response: from 1, I was fully aware, to 4, I was not aware at all); (5) ‘Was your state of mind intentional?’ (response: from 1, entirely intentional, to 4, entirely unintentional); (6) ‘How engaging were your thoughts?’ (response: from 1, not engaging, to 4, very engaging); (7) ‘How well do you think you have been performing?’ (response: from 1, not good, to 4, very good); (8) ‘How alert have you been?’ (response: extremely alert / alert / sleepy / extremely sleepy). Question 3 was displayed only if participants answered off-task in question 1. In this report, we focus only on questions (2) and (8).

##### Physiological Recordings and Preprocessing

The raw pupil size was corrected for the occurrence of blinks as in^2^. The timings of blinks were obtained through the EyeLink acquisition software. For each of these blinks, the pupil size was corrected by linearly interpolating the average signal preceding the blink onset ([-0.2, −0.1]s) and following the blink offset ([0.1, 0.2]s). The corrected signal was then low-pass filtered below 6Hz (two-pass Butterworth filter at the 4^th^ order). Finally, for blinks longer than 2s, data points between −0.1s prior to blink onset and 0.1s after blink onset were considered missing.

##### Detection of Slow Waves

In sleep, according to established guidelines^3^, only waves with peak-to-peak amplitude exceeding 75µV are defined as slow waves. In wakefulness, similar slow waves can be observed but with smaller amplitudes. Accordingly, previous studies relied on a relative rather than absolute threshold^4–6^. Here, we defined slow waves as the waves with absolute peak-to-peak amplitude within the top 10% of all the waves detected for a given EEG electrode and a given participant. On average, the detection threshold was 30 µV (average across N=26 participants and across all electrodes).

Figure 3a shows the average waveform of the slow waves detected on electrode Cz as well as the average waveform of waves detected during sleep recording in another published dataset (N=15 participants)^7^. To compute the average waveform of sleep slow waves, we applied the same algorithm to epochs of 20s scored as NREM2 and NREM3. Only slow waves with peak-to-peak amplitude over 75µV were considered.

We also compared the average waveform of slow waves with task-related Event Related Potentials (ERPs). To do so, we averaged the EEG signal time-locked to stimuli or response onsets at the electrode level. Figure S11a-c shows the average ERP across participants for electrode Cz as well as the scalp topographies of the voltage observed across the scalp at the time of the peak for electrode Cz. Finally, we also compared single-trial voltage values between the average waveforms for slow waves and stimulus-locked or response-locked ERPs (Figure S11d-f, data shown for one participant). Overall, these analyses showed a clear difference between the amplitude, waveform and topographies of slow waves compared to task-related ERPs.

#### Drift Diffusion Modelling

The Drift Diffusion Model (DDM) assumes that a decision variable noisily accumulates evidence from a starting point (*z*) with drift rate (*v*) towards one of two boundaries that represent choice alternatives (i.e. ‘Go’ or ‘No-go’; see Figure S7). The decision threshold (*a*) is the distance between the two boundaries and represents the amount of evidence that must be accumulated before a decision is made. Once the decision variable crosses a decision boundary, a response is made. Five parameters were fitted using a DDM approach: the participants’ initial bias for one of the two responses (decision bias, *z*), drift rates for Go and NoGo responses (*v_Go_* and *v_NoGo_*), the decision threshold (*a*), the non-decision time parameter (*t*). The last parameter, *t*, captures extra-decisional components, including stimulus encoding, response preparation and execution. We also extracted the drift rate bias (*v_Bias_*) as the difference between the absolute values of *v_Go_* (positive) and *v_NoGo_* (negative). We performed a model selection based on the Deviance Information Criteria (DIC), which assess goodness of fit while accounting for model complexity in hierarchical models^8^. With posterior predictive checks, we confirmed that our DDM was able to generate simulated behavioural data that are similar to the recorded data (Figure S8; based on the Fz model). For this check, we simulated 100 datasets based on the posterior distributions of HDDM parameters.

#### Statistics

A cluster-permutation approach (derived from ^9^) was applied to identify significant clusters in topographical maps. Candidate clusters were defined as neighbouring electrodes with a p-value below a threshold (called “cluster alpha”) of 0.025. For each candidate cluster, we computed the sum of the t-values for all the electrodes belonging to the cluster (which we will refer to as the “cluster statistics”). We then created permuted datasets by permuting the labels of the predictor within each subject, each task and each electrode (N=1,000 permutations). For each of these permuted datasets, we also identified the candidate cluster with maximal absolute cluster statistics. The cluster statistics from permutations formed a null distribution, against which we compared the cluster statistics from the real dataset. Clusters (real and permuted) with positive and negative cluster statistics were compared separately. A Monte-Carlo p-value was derived from this comparison (p_cluster_<0.05 means that a negative cluster has a cluster statistics below the 5^th^ percentile of the permuted distribution and that a positive cluster has a cluster statistics above the 95^th^ percentile of the permuted distribution). In cases where several cluster-permutations were performed in the same analysis (Fig. 5 and 6), we corrected the Monte-Carlo p-values of the real clusters with the Bonferroni method.

---

**Supplementary Table 1.**
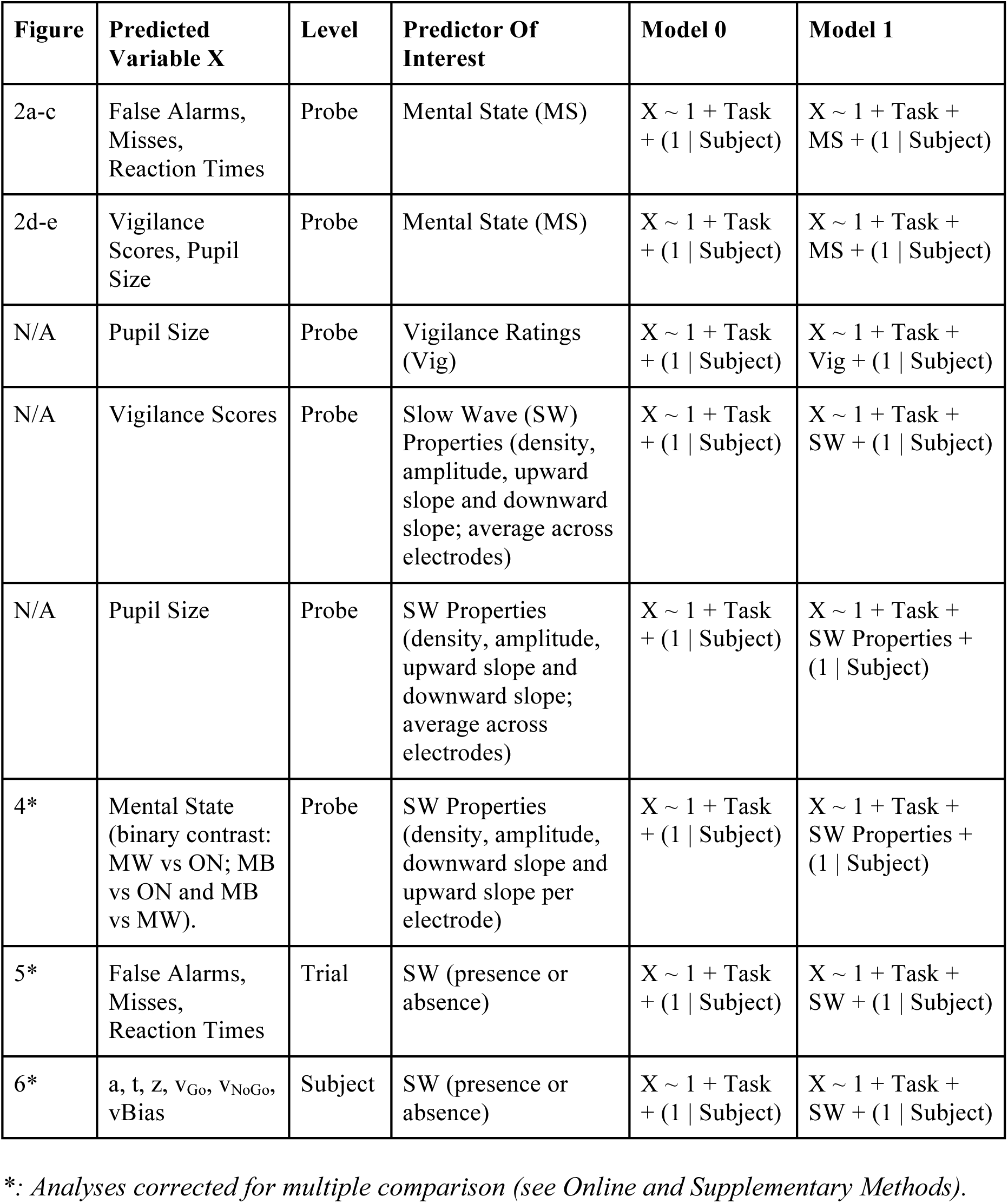
Summary of Linear Mixed-Effects Models.

**Supplementary Table 2.** Effect of normalisation on behavioural analyses. The analyses of the effect of mental states (model comparisons and post-hoc contrasts) were consistent even after normalizing Misses, False alarms and Reactions Times (RT) for each participant by the average obtained on ON probes (“Normalised” column). The “State” row shows the result of the likelihood ratio test (chi-squared value). Stars denote the corrected significance level (Bonferroni correction for 6 likelihood ratio tests: ***: p<0.001, **: p<0.01, *: p<0.05). The estimates of the post-hoc contrasts (MW vs ON, MB vs ON, MB vs MW) obtained from the mixed effect models are also reported, with the corresponding 95% confidence intervals. Bold fonts signal the significant results (p<0.05 for model comparisons, or confidence intervals excluding 0 for the post-hoc contrasts).

|  |  | Normalised | Before Normalisation |
| --- | --- | --- | --- |
| Misses (%)<br>in Go trials | State | $\chi^2(2)=33.4$ (***) | $\chi^2(2)=36.0$ (***) |
|  | *MW vs ON* | -0.0096 [-0.015, -0.0041] | -0.011 [-0.016, -0.0052] |
|  | *MB vs ON* | -0.023 [-0.032, -0.015] | -0.023 [-0.032, -0.015] |
|  | *MB vs MW* | -0.014 [-0.022, -0.0055] | -0.013 [-0.021, -0.0047] |
| False<br>alarms (%)<br>in NoGo<br>trials | State | $\chi^2(2)=88.5$ (***) | $\chi^2(2)=115.9$ (***) |
|  | *MW vs ON* | -0.24 [-0.29, -0.19] | -0.20 [-0.24, -0.17] |
|  | *MB vs ON* | -0.20 [-0.26, -0.19] | -0.17 [-0.23, -0.12] |
|  | *MB vs MW* | 0.064 [-0.014, 0.14] | 0.028 [-0.028, 0.084] |
| RT (s) in<br>Go trials | State | $\chi^2(2)=33.4$ (***) | $\chi^2(2)=16.9$ (**) |
|  | *MW vs ON* | -0.0005 [-0.013, 0.011] | -0.0025 [-0.0096, 0.0045] |
|  | *MB vs ON* | **0.039 [0.022, 0.057]** | **0.019 [0.0088, 0.030]** |
|  | *MB vs MW* | **0.040 [0.022, 0.058]** | **0.022 [0.011, 0.032]** |

**Supplementary Table 3.**
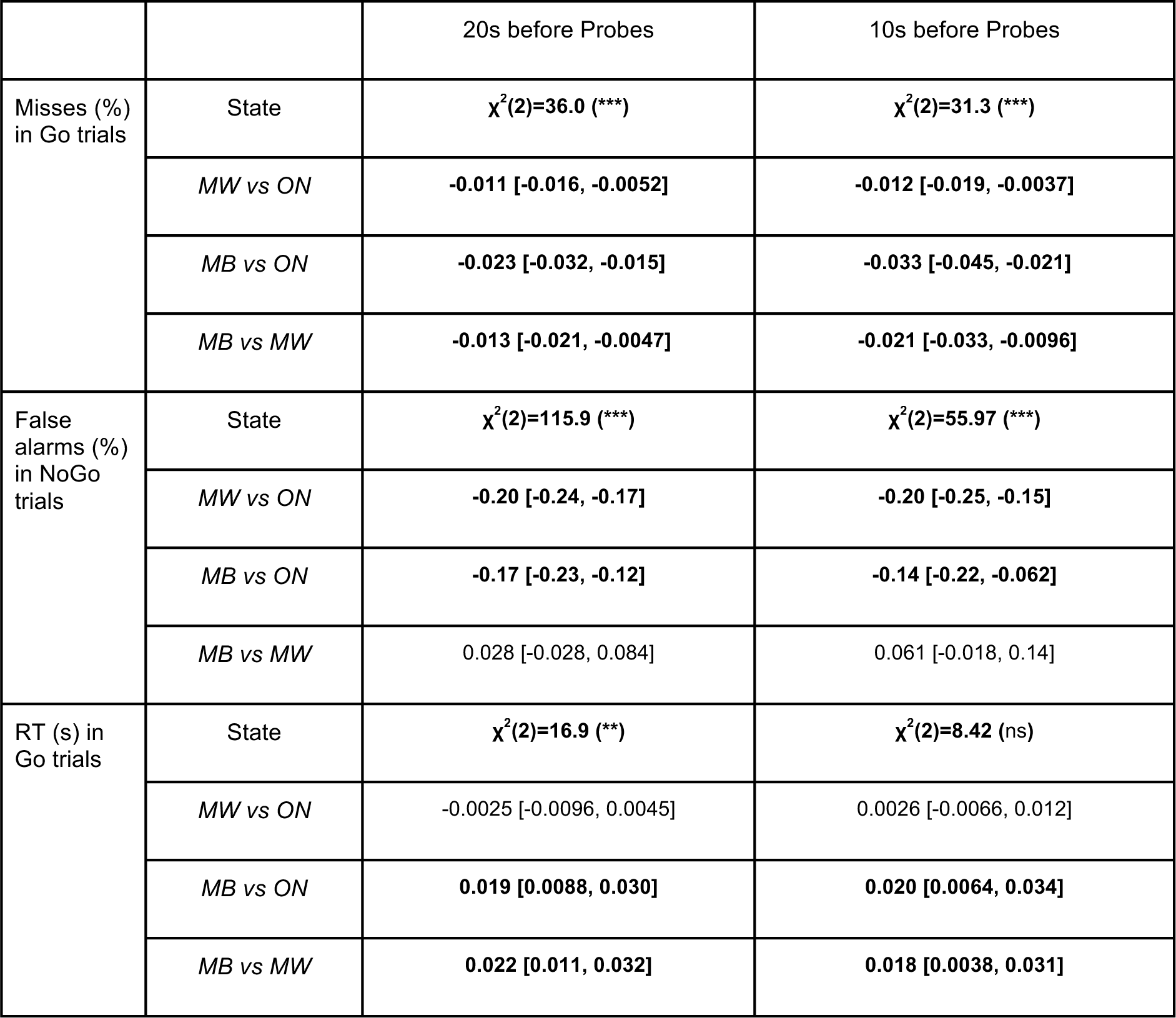
Effect of different pre-probe window sizes on behavioural analyses. The analyses of the effect of mental states (model comparisons and post-hoc contrasts) were replicated using a shorter window of 10s (right column) instead of 20s before probe onsets (left column). Likelihood Ratio Tests are reported for the effect of State on the correctness for Go and NoGo trials as well as the Reaction Times (RT) for Go Trials. Stars denote the corrected significance level (Bonferroni correction for 6 comparisons: ***: p<0.001, **: p<0.01, *: p<0.05). The estimates for the post-hoc contrasts and the corresponding 95% confidence intervals are also shown. Bold fonts signal the significant results (p<0.05 for model comparisons, or confidence intervals excluding 0 for the post-hoc contrasts).

**Supplementary Figure 1.**
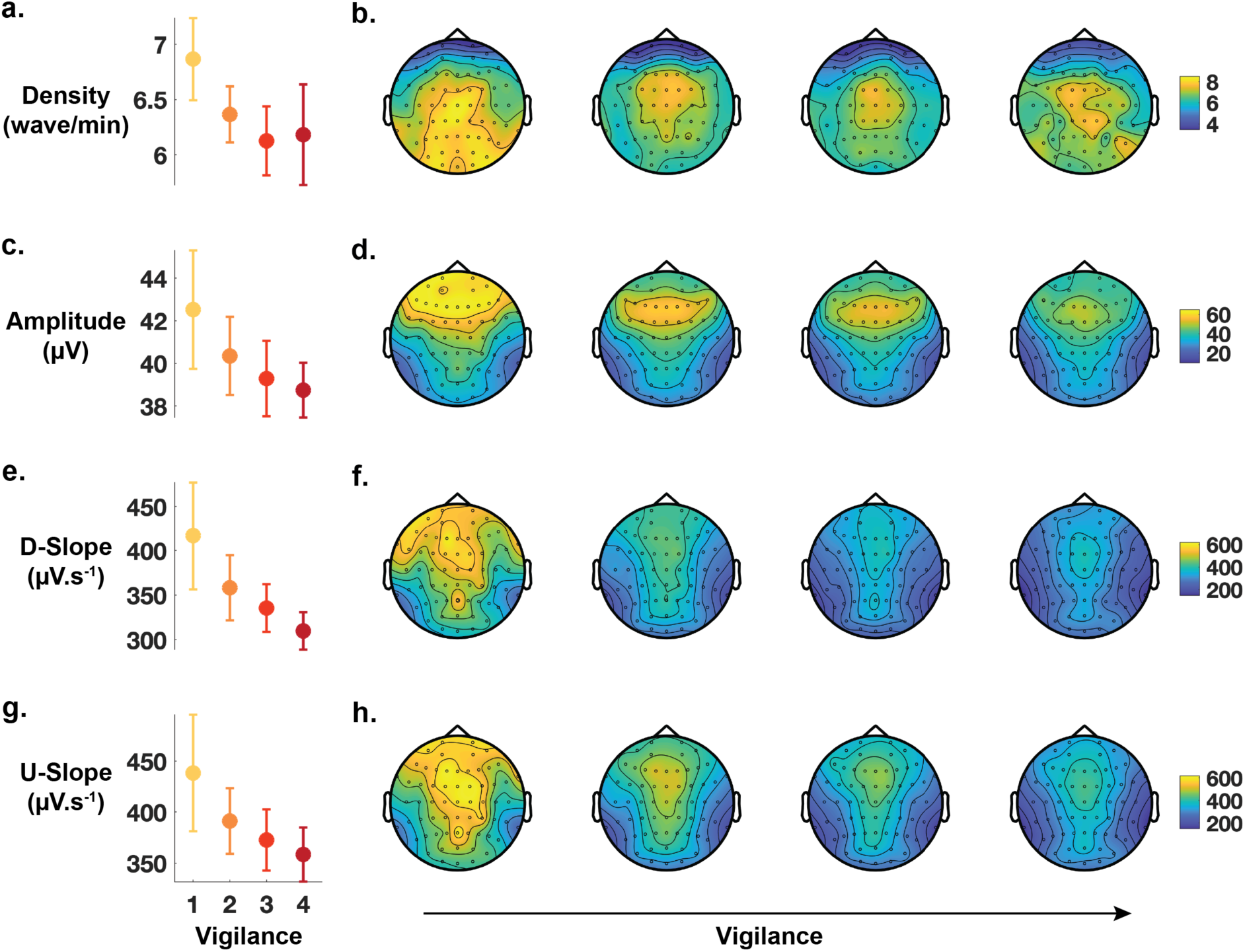
Sleepiness is associated with more, larger and steeper slow waves. Slow waves detected within 20s of probes’ onset were separated according to participants’ vigilance ratings (from 1 (extremely sleepy) to 4 (extremely alert)) provided after each probe. The density (a), amplitude (c), downward slope (D-Slope, e) and upward slope (U-Slope, f), averaged across participants and electrodes, are shown with error-bars showing the standard-error-of-the-mean across participants. Topographies of the average density (b), amplitude (d), downward slope (f) and upward slope (h) are also shown for the different vigilance levels.

**Supplementary Figure 2.**
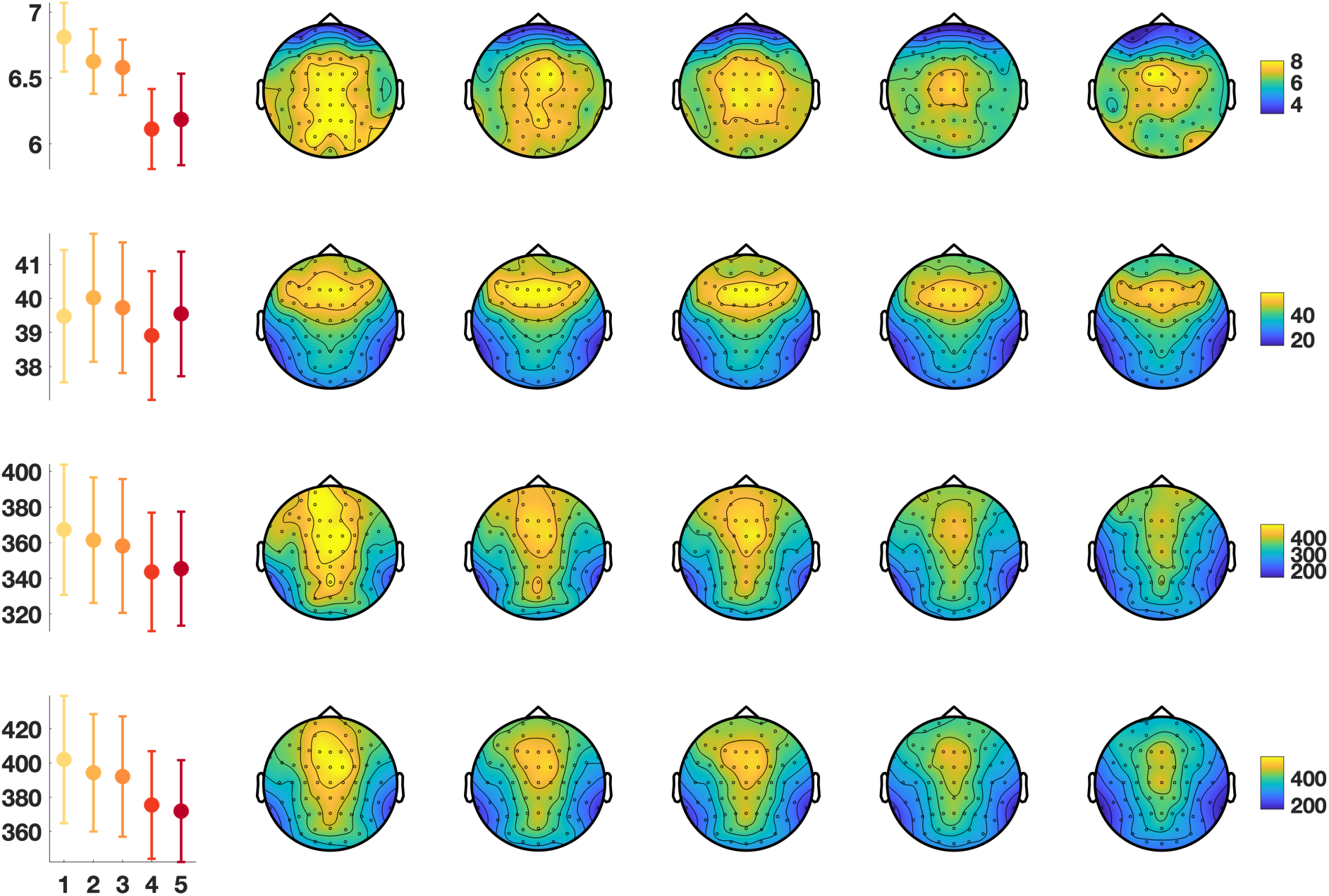
A reduction in pupil size is associated with more, larger and steeper slow waves. Slow waves detected within 20s of probes’ onset were separated according to participants’ pupil size (binned, see Methods) computed on the same window. The density (a), amplitude (c), downward slope (D-Slope, e) and upward slope (U-Slope, f), averaged across participants and electrodes, are shown with error-bars showing the standard-error-of-the-mean across participants. Topographies of the average density (b), amplitude (d), downward slope (f) and upward slope (h) are also shown for the different pupil size bins.

**Supplementary Figure 3.**
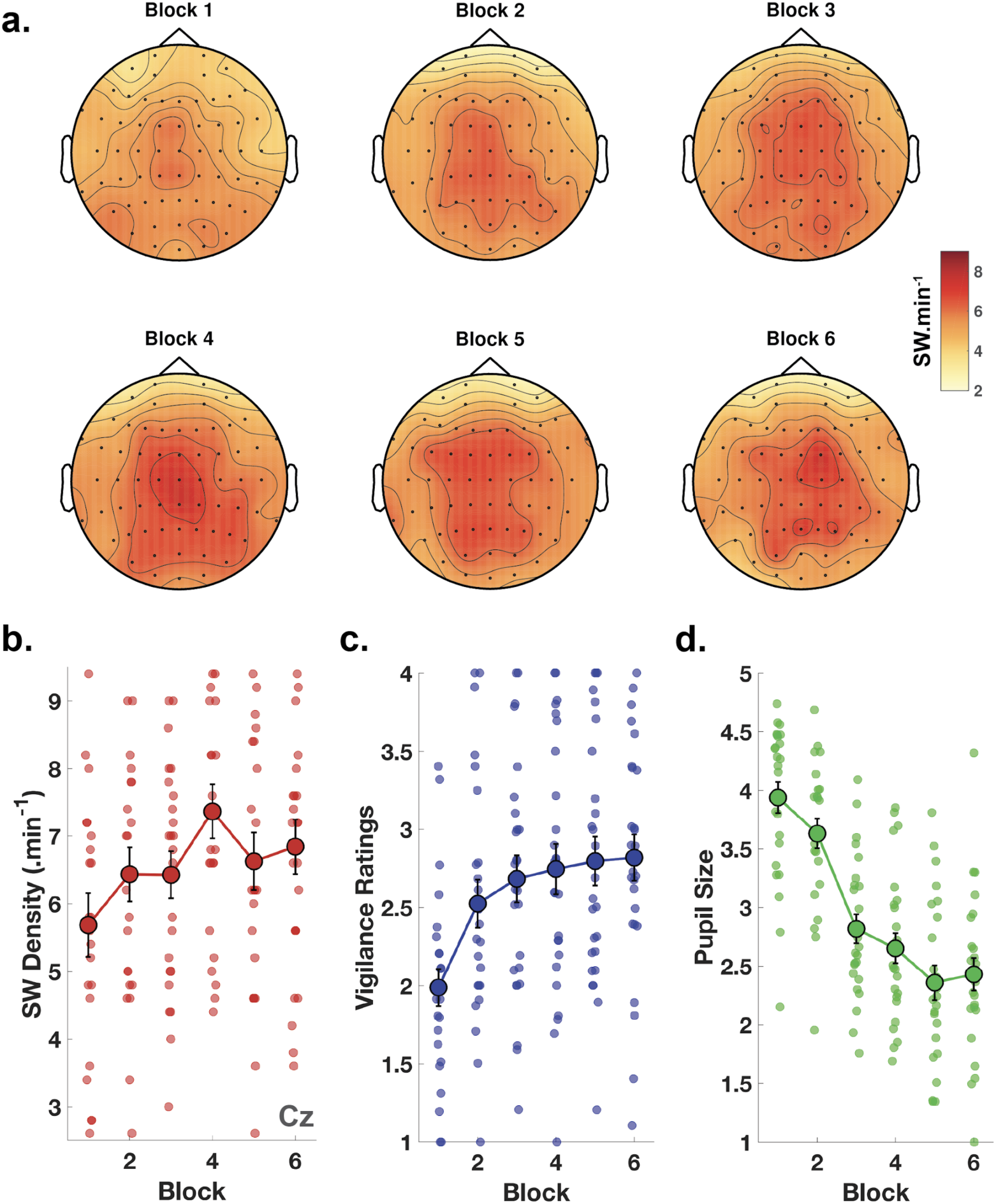
Slow waves increase with time spent on task. (a) Topographies of the temporal density of slow waves (slow waves per minute, averaged across all 26 subjects). An increase in the number of slow waves, maximal over central electrodes, can be seen from the first to the last experimental block. (b-d) Slow wave density for electrode Cz (b), vigilance ratings (c) and pupil size (d) averaged within each block. Connected larger dots show the average across participants (error-bars: SEM). Individual semi-transparent circles show individual subjects.

**Supplementary Figure 4.**
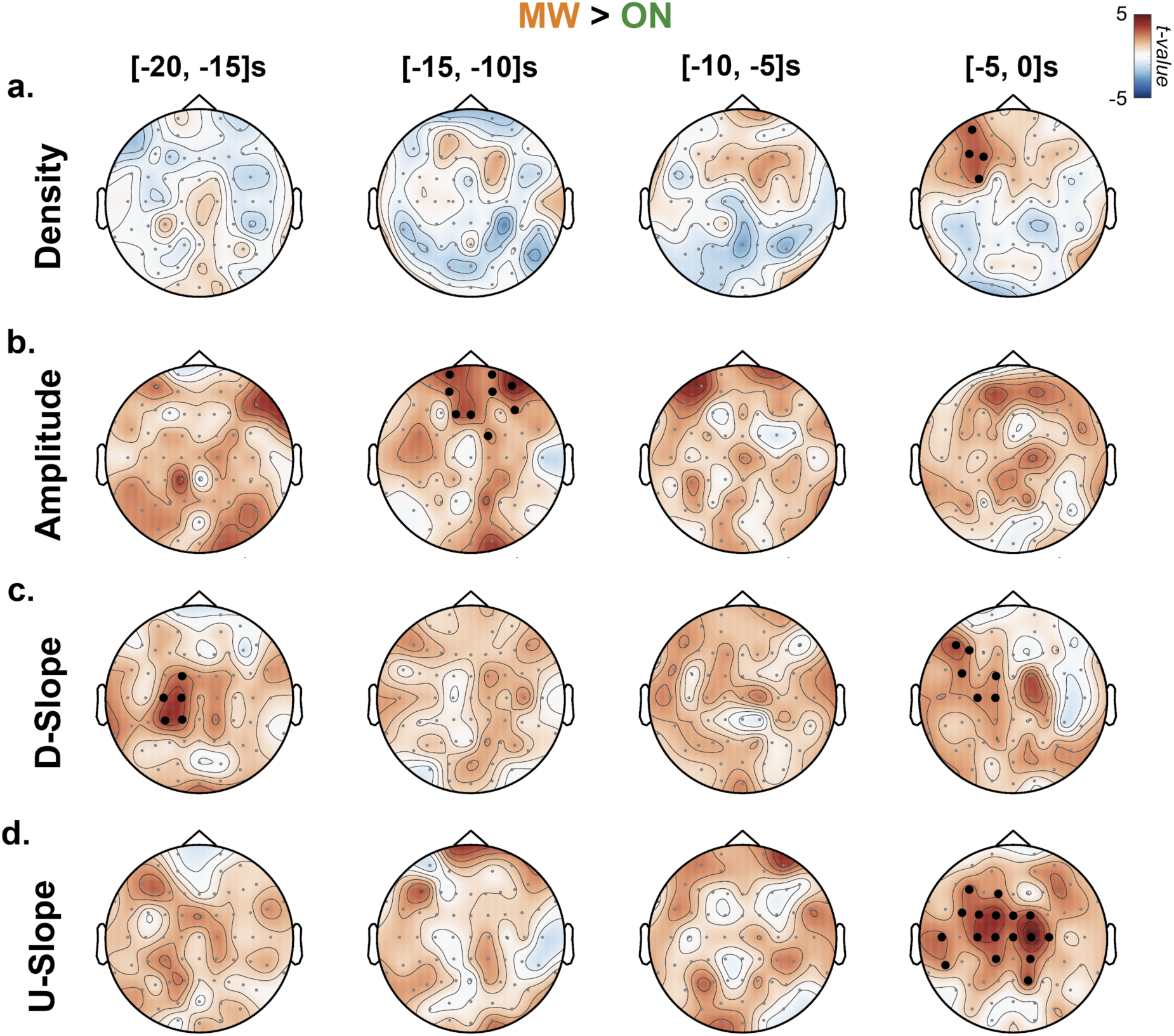
Spatiotemporal dynamics of the effect of slow-waves properties on mental states (MW vs ON). Mixed-Effects Models were used to quantify the impact of slow-wave properties (**a**: Density; **b**: Amplitude; **c**: Downward Slope (D-Slope); **d**: Upward Slope (U-Slope)) on mental states as in Figure 4. Slow waves were detected on 4 different windows: [-20, −15]s, [-15, −10]s, [-10, −5]s and [-5, 0]s before probe onsets. Slow-wave properties were extracted for each electrode and used to predict the MW vs ON contrast. Topographies show the scalp distribution of the t-values associated with each slow-wave parameter and electrode. Black dots denote significant clusters of electrodes (p_cluster_<0.05 corrected for 48 comparisons (Figure S4-6) using a Bonferroni approach).

**Supplementary Figure 5.**
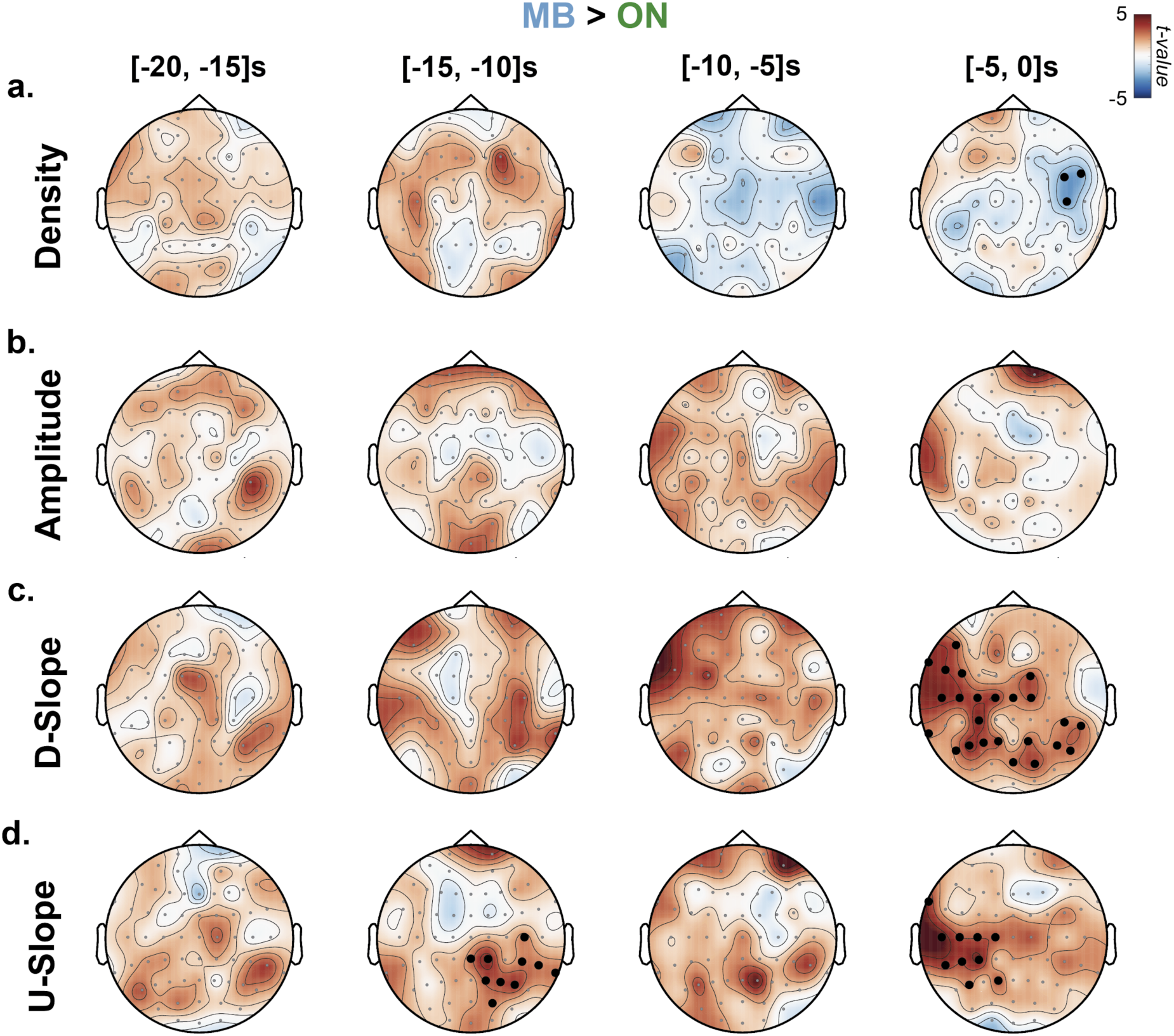
Spatiotemporal dynamics of the effect of slow-waves properties on mental states (MB vs ON). Same format as in Supplementary Figure 4.

**Supplementary Figure 6.**
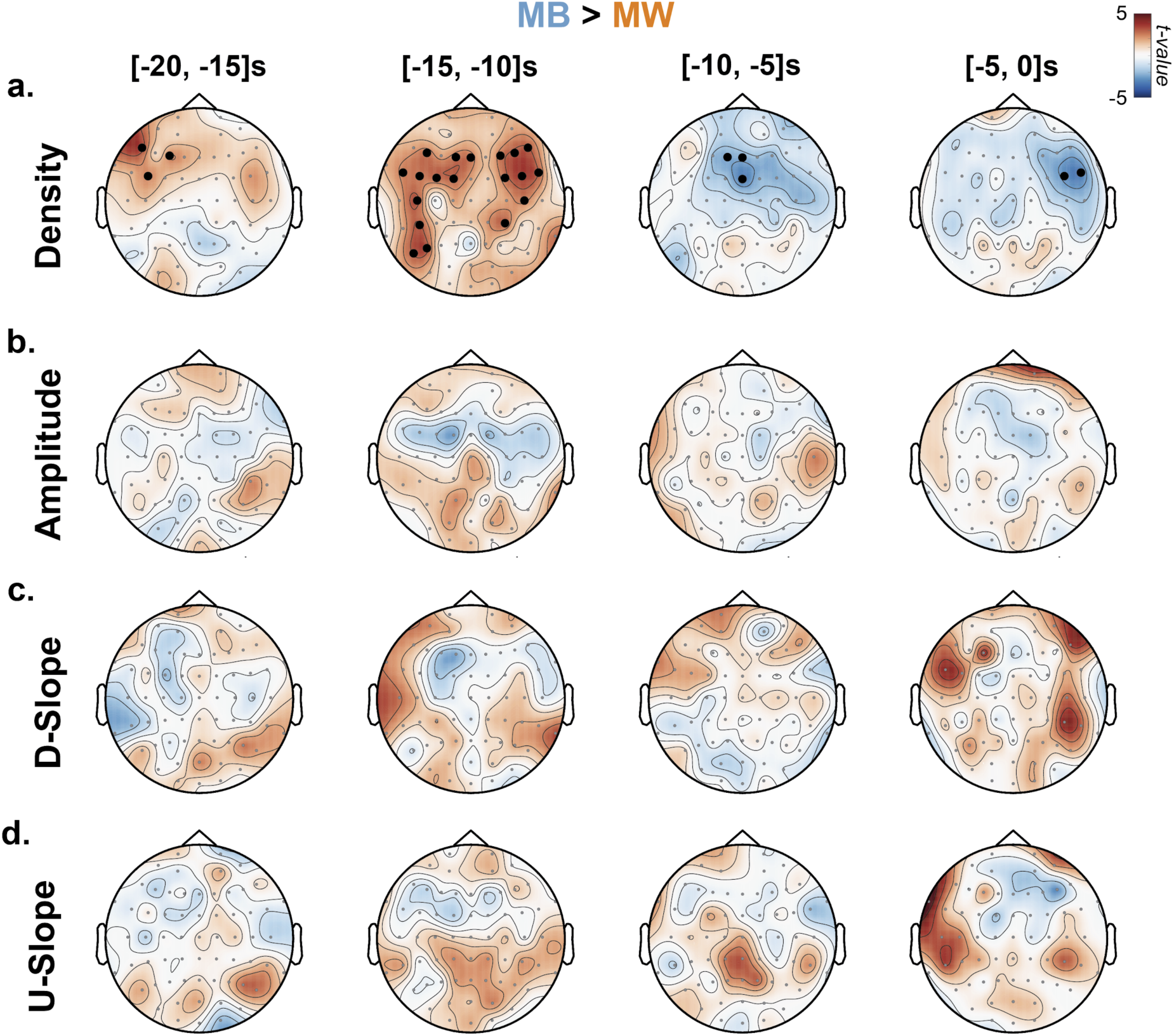
Spatiotemporal dynamics of the effect of slow-waves properties on mental states (MB vs MW). Same format as in Supplementary Figure 4.

**Supplementary Figure 7.**
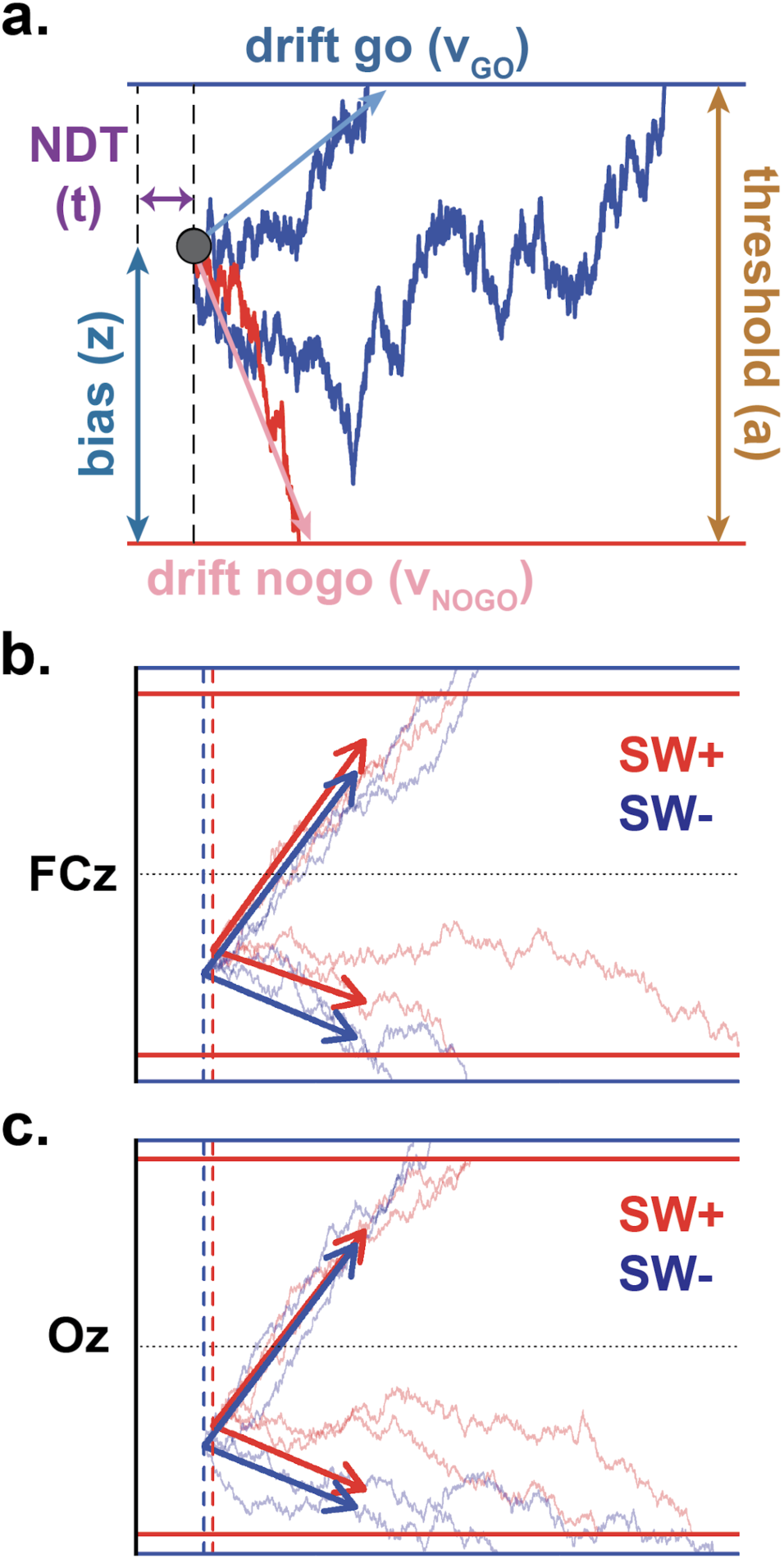
Drift Diffusion Modelling. **a:** The Go/NoGo tasks were modelled according to the Drift Diffusion Model (DDM, see Online and Supplementary Methods). The following parameters were estimated: threshold (a), non-decision time or NDT (t), bias (z), drift rate for Go trials (v_Go_), drift rate for NoGo trials (v_NoGo_). We then also computed the drift bias: abs(v_Go_)-abs(v_NoGo_). The figure shows a graphical representation of these parameters. Note that here, drift rates for NoGo trials are negative. **b-c:** Graphical representation of decision processes using the parameters obtained by the DDM for trials with (SW+) or without (SW-) slow waves. The presence or absence of slow waves was determined for each electrode (FCz (b; frontal) and Oz (c; posterior)). Note the reduction in decision threshold, drift rates and bias associated with slow waves but the increase in NDT.

**Supplementary Figure 8.**
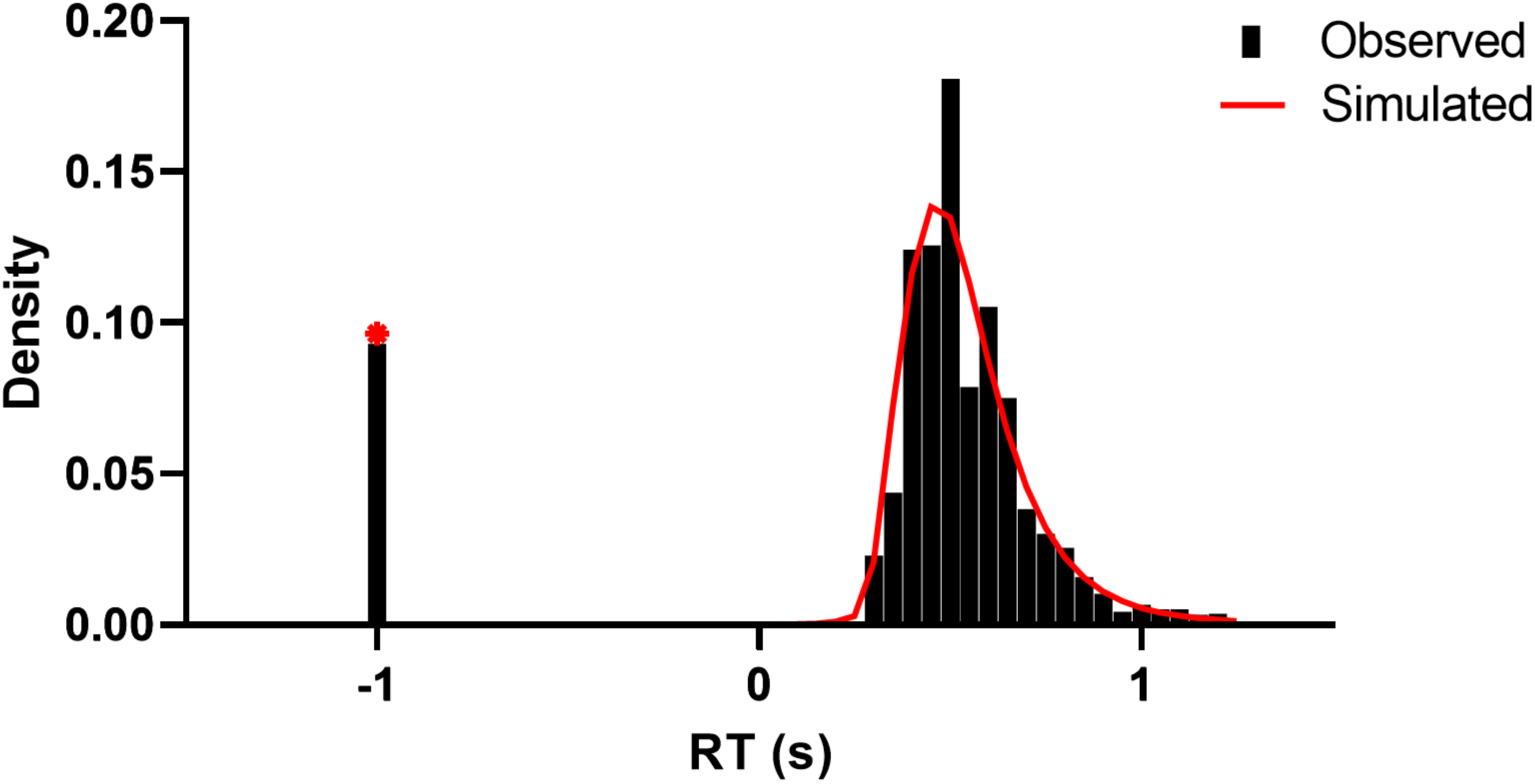
HDDM Fit to Behavioural Data. Posterior predictive checks of Go/No-Go DDM fit to the behavioural data. Observed data for all participants (black bars) are plotted underneath model-predicted RT distributions and No-Go choice proportions (red lines). Positive distribution represents the normalised frequency of reaction times (RT) from Go responses. Negative bin at RT=-1 represents the proportion of No-Go responses.

**Supplementary Figure 9.**
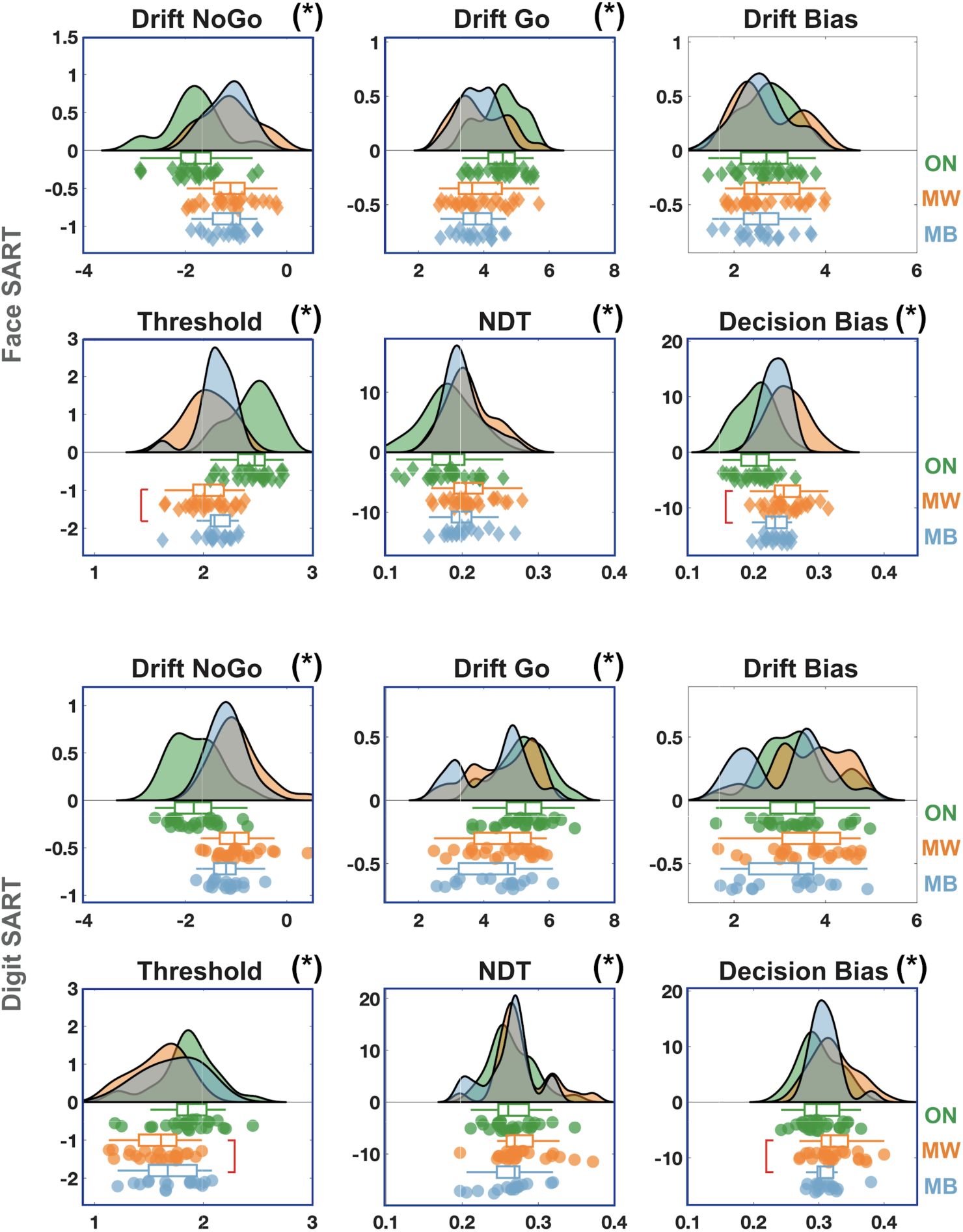
Impact of mental states on HDDM parameters. Hierarchical Drift Diffusion Modeling (HDDM, see Methods) was applied to the Reaction Times obtained in the Face (top) and Digit (bottom) SART. The parameters were fitted for each task and mental state. Each panel shows the distribution of the estimated variable for individual participants (drift for NoGo and Go trials, drift bias, threshold, non-decision time (NDT) and decision bias; see Methods). For each plot, coloured areas show the smoothed distribution of individual data points (see Methods). Diamonds and circles show individual estimates for the Face and Digit SART respectively. Box plots show the 1^st^ and 3^rd^ quartiles (edges) as well as the median (middle bar). Blue boxes around individual plots and stars next to the titles indicate variables with significant state-effects (model comparison, see Methods; *: p<0.05, Bonferroni correction for 12 comparisons). Significant differences between MW and MB (for threshold and decision bias) are highlighted with red brackets.

**Supplementary Figure 10.**
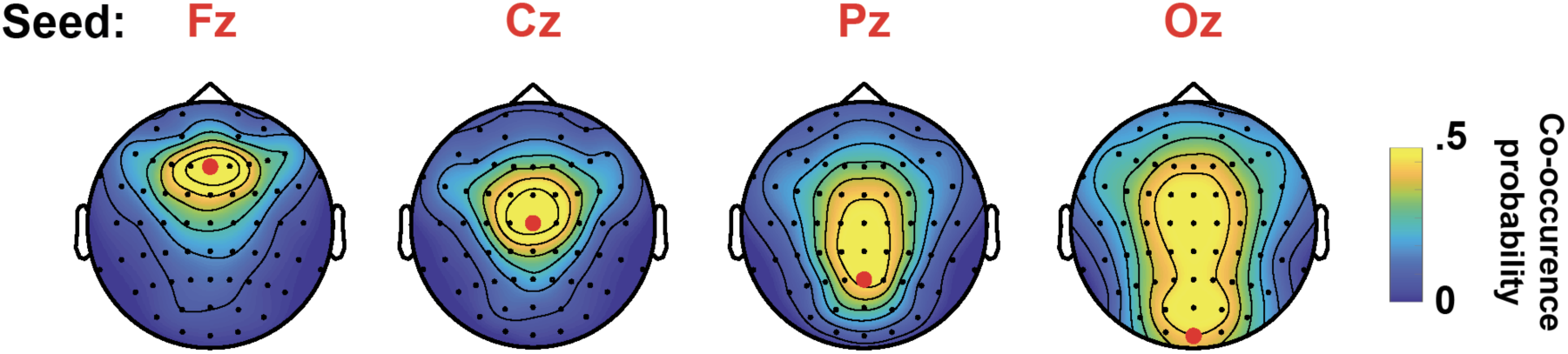
Spatial expanse of slow waves. Four seeds electrodes were selected along the scalp midline from the front (Fz) to back (Oz). For each slow wave detected in these seed electrodes, we computed the probability that slow waves were also observed in the other electrodes. The average co-occurrence probability averaged across participants (N=26) is shown for each seed electrode. Note that slow waves detected over Fz tend to co-occur with other slow waves only in a limited number of neighboring frontal electrodes whereas occipital slow waves (Oz) tend to co-occur with other slow waves in both frontal and posterior electrodes (more widespread).

**Supplementary Figure 11.**
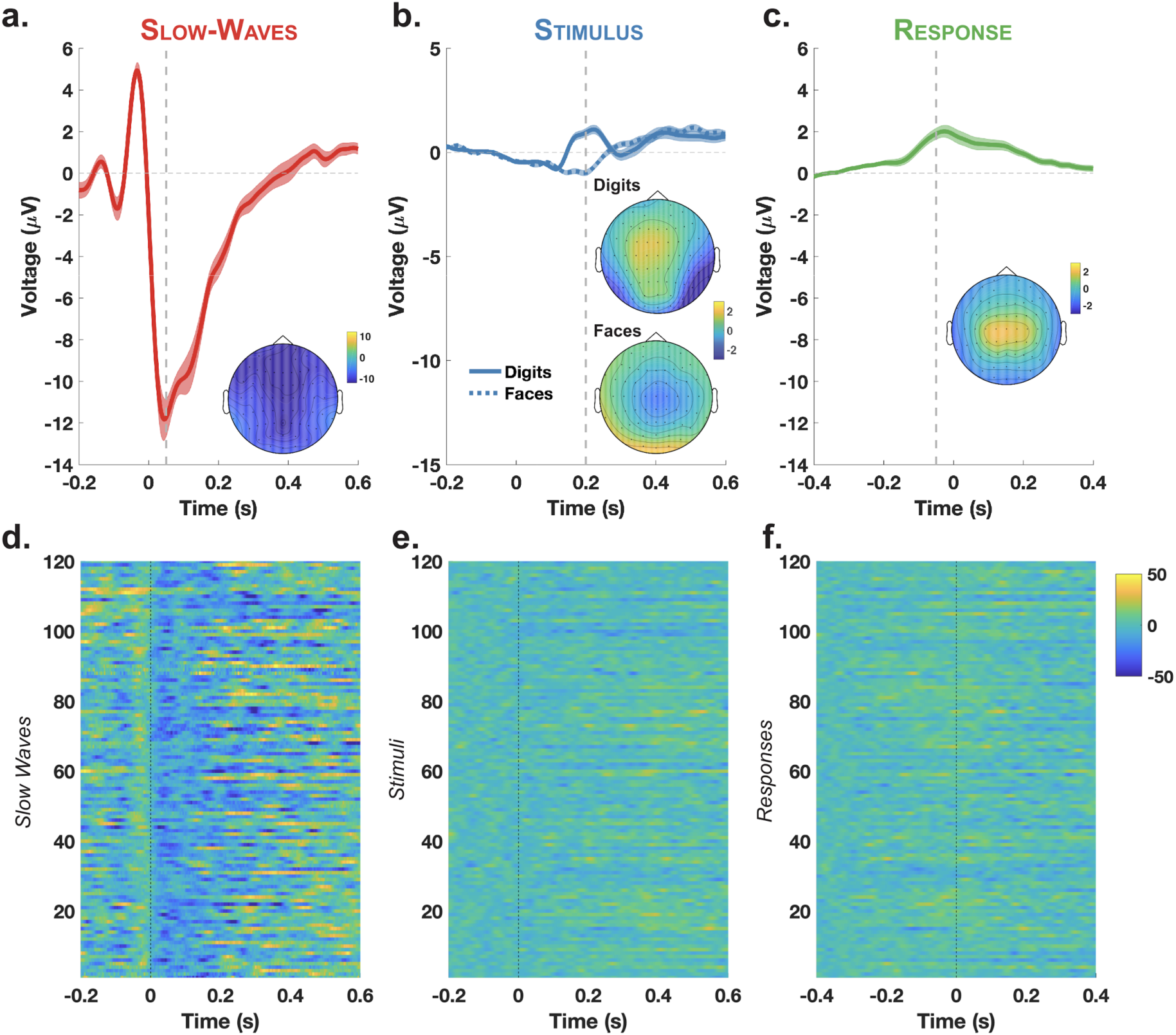
Slow waves compared to stimulus and response-locked activity. Event-related potentials averaged over electrode Cz and across participants for slow waves (a), stimulus-locked activity (b) and response-locked activity (c). Slow waves’ EEG time courses are aligned with the first negative crossing before the negative peak. Stimulus-locked responses are aligned on the onset of the digit (full line) or face (dotted line) stimuli. Response-locked responses are aligned to the onset of participants’ responses. Shaded areas show the SEM across participants (N=26). Insets show the topography of the average voltage at the times shown by the vertical dotted lines (a: 0.05s; b: 0.2s; c: −0.05s), which approximates the absolute maximum of each ERP. (d-f) Voltage of the event-related potentials per trial for one participant. Each row corresponds to (d) an individual slow wave, (e) a stimulus presentation (face or digit), (f) a motor response. We limited the number of rows in (e) and (f) to match the number of detected slow waves (N=120) in (d).

**Supplementary Figure 12.**
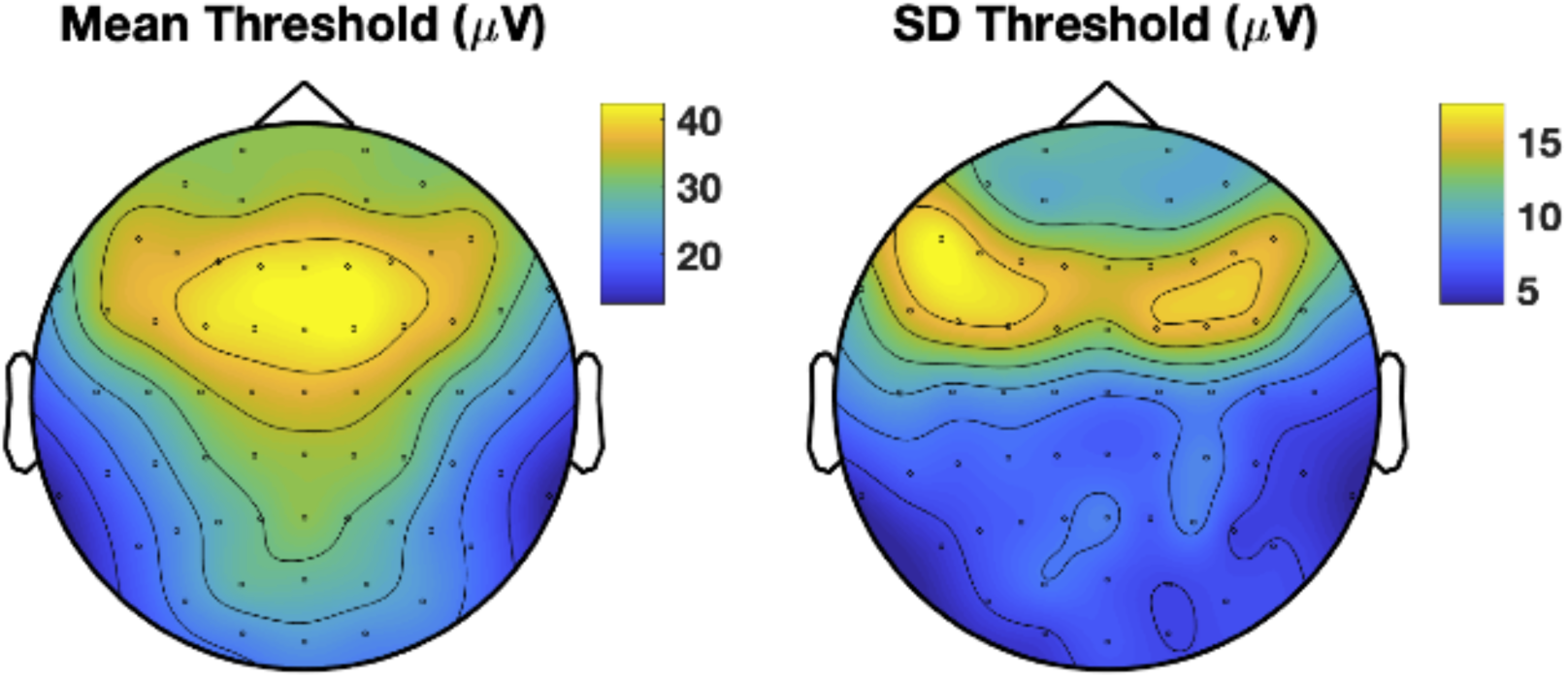
Scalp topography of the average and standard-deviation of the slow-wave detection threshold. Only slow waves whose amplitude is within the top-10% of the distribution for each electrode and participant were analysed. The left topography shows the value, in voltage, of this threshold for each electrode, averaged across participants. The right topography shows the standard deviation (SD) of the threshold values across participants.

